# Metabolic control of immune-competency by odors in *Drosophila*

**DOI:** 10.1101/718056

**Authors:** Sukanya Madhwal, Mingyu Shin, Manish K Joshi, Ankita Kapoor, Pirzada Mujeeb Ur Rehman, Kavan Gor, Jiwon Shim, Tina Mukherjee

**Affiliations:** Institute for Stem Cell Science and Regenerative Medicine (inStem), Bellary Road, Bangalore 560065, India; Department of Life Science, College of Natural Science, Hanyang University, Seoul, 04763, South Korea; Vellore Institute of Technology, Katpadi Road, Vellore, Tamil Nadu 632014, India

**Author notes:** Manipal Academy of Higher Education, Manipal, Karnataka 576104, India (Sukanya Madhwal, Ankita Kapoor). Aix Marseille Université, CNRS, Institut de Biologie du Développement de Marseille (IBDM), Marseille, France. University of Cologne, CECAD-Cluster of Excellence, Joseph-Stelzmann-Str. 26, Köln 50931, Germany (Pirzada Mujeeb Ur Rehman).

**Keywords:** Olfaction, Hematopoiesis, GABA-shunt, Succinate, HIFα, Metabolism, Immunity, Infection

## Abstract

*Drosophila* blood-progenitor cells generate an inflammatory cell-type termed lamellocyte, in response to parasitic wasp-infections. In this study we show that olfaction primes lamellocyte potential. Specifically, larval odor-detection mediated release of systemic γ-aminobutyric acid (GABA) from neurosecretory cells, is detected and internalized by blood progenitor-cells. GABA catabolism through the GABA-shunt pathway prevents Sima (HIFα) protein degradation. Sima is necessary and sufficient for lamellocyte induction. However, limited systemic GABA availability during development restricts blood-progenitor Sima levels and consequently their lamellocyte potential. Preconditioning *Drosophila* larvae in odor environments mimicking parasitoid-threatened conditions raises systemic GABA and blood-progenitor Sima levels. As a result, infection responses in these animals are rapid and efficient. Overall, this study explores the importance of sensory control of myeloid-immunity and unravels the adaptive influence of environmental odor-experience on myeloid-metabolism and priming innate-immune potential.

## INTRODUCTION

Hematopoiesis in *Drosophila* gives rise to three blood cell types: plasmatocytes, crystal cells and lamellocytes, with characteristics that are reminiscent of the vertebrate myeloid lineage. Of these, lamellocytes, which are undetectable in healthy animals, appear upon infections with the parasitic wasp, *Leptopilina boulardi* that triggers their development(Crozatier et al., 2004). Within a few hours of wasp-egg deposition, the *Drosophila* larval hematopoietic system activates a series of cellular innate immune responses leading to massive differentiation of blood cells into lamellocytes. This includes trans-differentiation of circulating and sessile plasmatocytes and differentiation of multipotent blood-progenitor cells of the larval hematopoietic organ termed the “lymph gland” (Anderl et al., 2016; Honti et al., 2010; Markus et al., 2009; Stofanko et al., 2010). As lymph gland progenitor cells differentiate, the gland ultimately disintegrates to release its blood cells into circulation(Lanot et al., 2001) necessary for successful neutralization of the deposited wasp-egg (Louradour et al., 2017). While the precise contributions of the two hematopoietic pools, both circulating and lymph gland, remains an active area of investigation, their combined response contributes towards robust lamellocyte numbers reaching a maximum at 48 hours after wasp-egg laying(Lanot et al., 2001). Characterized by their large flattened appearance, lamellocytes encapsulate the deposited wasp-eggs and melanize them, facilitating their effective clearance(Rizki and Rizki, 1992). Blood cells therefore maintain a on demand – adapted hematopoietic process to develop lamellocytes. This innate competitiveness provides a defence mechanism for the fly to limit parasitoid success. Developmental programs that establish lamellocyte potential and capacitate the hematopoietic system to respond when in need, forms the central focus of our investigation.

Lymph gland blood-progenitor cells rely on systemic cues of olfactory origin for their maintenance(Shim et al., 2013). The *Drosophila* larval olfactory system contains 25 specific odorant receptors (OR) in 21 olfactory receptor neurons (ORNs)(Vosshall and Stocker, 2007). Orco (Or83b) is an atypical odorant receptor protein, expressed in every ORN and is necessary to respond to all odors(Larsson et al., 2004). Odors are sensed by the larval dorsal organ, which is innervated by dendrites of these ORNs that project to specific glomerulus of the larval antennal lobe. Here, ORNs form excitatory synapses with projection neurons (PNs) whose axons innervate into regions of the brain representative of higher order information processing. The different glomeruli are interconnected by excitatory or inhibitory local interneurons that fine-tune the ORN-PN network(Masse et al., 2009).

Earlier work has demonstrated a dependence on olfaction induced systemic γ-aminobutyric acid (GABA) availability in the maintenance of blood progenitor cells. Specifically, during *Drosophila* larval development, olfaction stimulates the release of GABA from neurosecretory cells of the brain which leads to systemic activation of GABA_B_R signaling of the lymph gland blood progenitor cells. The resulting GABA signal causes elevation of cytosolic calcium, which is necessary to support progenitor maintenance(Shim et al., 2013). This study established the mechanistic basis for odor induced control of progenitor homeostasis. Cross-talk between olfaction and the hematopoietic system is also reported in higher animals(Strous and Shoenfeld, 2006) and is not only limited to *Drosophila*. However, any physiological relevance for this communication remains obscure.

In this study, we demonstrate a role for the larval olfactory system in priming cellular immune potential that is necessary to sustain a demand-adapted immune response via co-opting the use of systemic GABA as a metabolite. The neuronally released GABA derived upon olfactory stimulation, is internalized by blood cells via Gat, the GABA-transporter. Intracellularly, GABA is catabolized through the GABA-shunt pathway to generate succinate, which is necessary for the stabilization of Sima, orthologous to the mammalian HIFα (Hypoxia inducible factor α) protein, a well-characterized transcription factor generally implicated in hypoxia response(Lavista-Llanos et al., 2002). Resulting Sima in immune cells is necessary and sufficient to capacitate blood cells with the ability to differentiate into lamellocytes. Further explorations to delineate the physiological relevance of this long-range cross-talk in controlling lamellocyte-competency unravels a sensory modality of pre-conditioning immune-competency and efficient lamellocyte formation. Thus showing that, odor detection in *Drosophila* establishes a long-range metabolic axis to control basic immune-competency during blood development.

## RESULTS

### Intracellular GABA regulates hematopoietic progenitor immune-competency

We first investigated whether GABA levels in blood cells is responsive to *L. boulardi* mediated wasp-immune challenge. To detect this, we undertook immuno-staining with anti-GABA antibody in lymph gland samples obtained from larvae 6-hours post immune challenge. In comparison to intracellular GABA (iGABA) levels detected in blood cells in normal homeostatic conditions (Fig. 1a), around a 2-3 fold increase in iGABA expression was detected post wasp-infection (Fig. 1b, c and and Supplemental Fig. 1a, a’ see Methods for iGABA staining protocol). The increased iGABA levels detected in blood cells following the immune-challenge led us to investigate the consequences of manipulating GABA levels.

**Figure 1.**
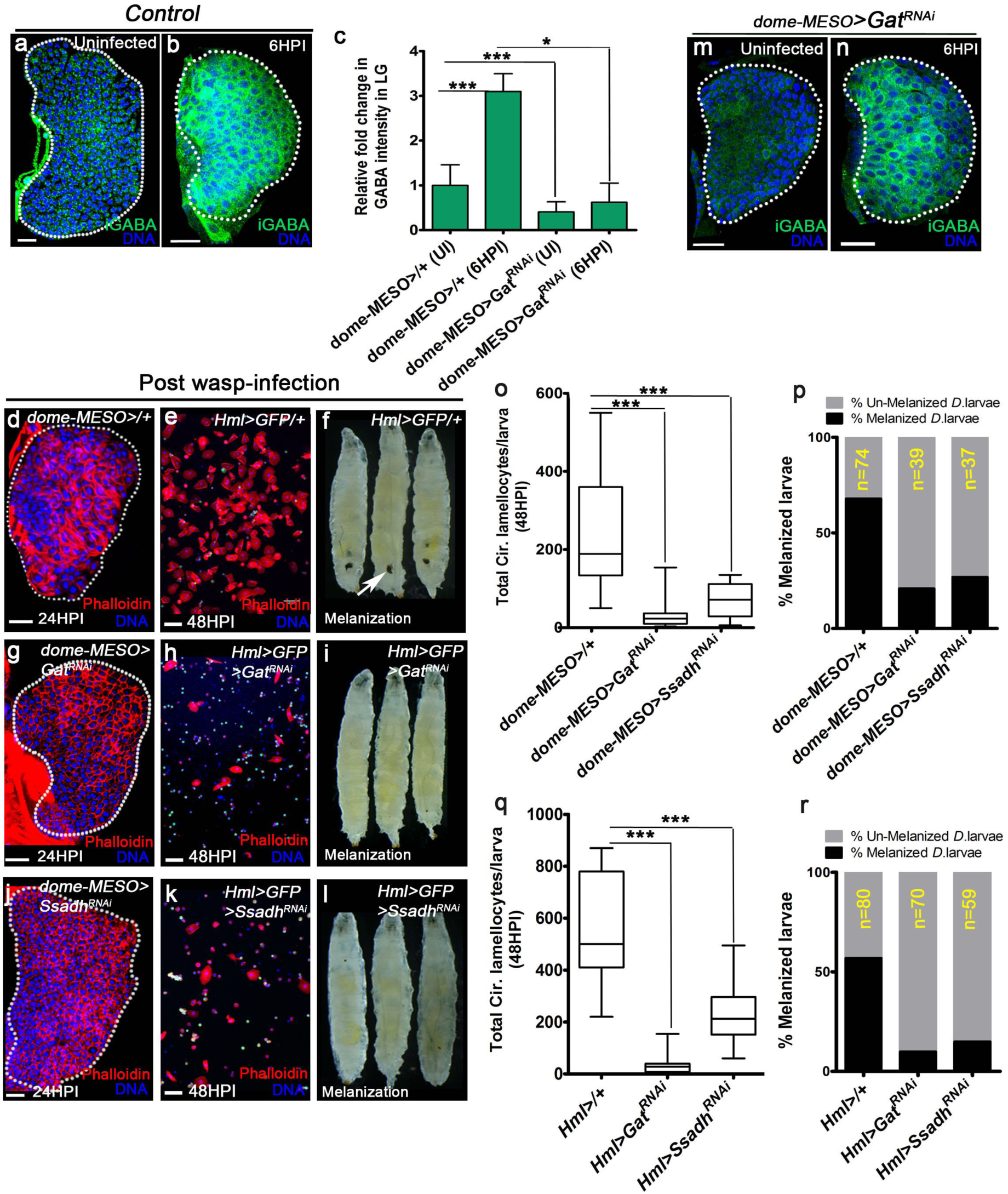
GABA-uptake and catabolism is necessary for lamellocyte formation. DNA is marked with DAPI (blue), intra-cellular GABA (iGABA, green) and blood cells are marked with phalloidin (red). Lamellocytes are discriminated by their characteristic large flattened morphology. In panels **e, h** and **k** scale bars = 50μm and **a, b, d, g, j, m, n**= 20μm. UI indicates un-infected, HPI indicates <u>h</u>ours <u>p</u>ost wasp-infection. In panels **c**, bar is Standard error of mean (SEM) and in **o, q** median is shown in the box plots and vertical bars represent upper and lowest cell counts. Statistical analysis, un-paired *t*-test, two-tailed in panel **c** and Mann-Whitney test, two-tailed in panels **o** and **q**. “n” is total number of animals analyzed. (**a, b**) Control (*domeMESO>GFP/+*) lymph gland from (**a**) un-infected *Drosophila* larvae show punctated iGABA staining in all blood cells (anti-GABA antibody staining in 1X PBS + 0.3%Triton X-100). In comparison to (**a**), iGABA levels detected in (**b**) lymph glands at 6HPI is elevated. See corresponding quantifications in **c**. (**c**) Quantifications of relative fold change in iGABA intensities. Control UI (*domeMESO>GFP/+*, n=10), Control 6HPI (*Dome>GFP/+*, n=11, ***p<0.0001), *domeMESO>Gat^RNAi^* UI (n= 10, ***p<0.0001) and *domeMESO>Gat^RNAi^* 6HPI (n= 13, *p=0.047). Bars represent standard error of mean (SEM). See Supplemental Fig. 1a, f and g for mean iGABA intensities. (**d-f**) Wasp-infection response in controls showing lamellocytes in **(d)** lymph gland, (**e**) circulation and (**f**) melanization of the deposited wasp-egg (black melanotic capsules, marked with white arrow). (**g**) Expressing *Gat^RNAi^* in progenitor cells (*domeMESO-Gal4, UAS-GFP; UAS-Gat^RNAi^*) causes reduction in lamellocyte numbers both in the lymph gland (see quantification in Supplemental Fig. 1b) and circulation (quantifications in **o**) and melanization response (quantifications in **p**). (**h, i**) Expressing *Gat^RNAi^* in differentiating hemocytes (*Hml^Δ^-Gal4, UAS-GFP; UAS-Gat^RNAi^*) leads to reduction in (**h**) circulating lamellocytes (quantifications in **q**) and (**i**) melanization response (quantifications in **r**). (**j**) Expressing *Ssadh^RNAi^* in progenitor cells (*domeMESO-Gal4,, UAS-GFP; UAS-Ssadh^RNAi^*) causes reduction in lamellocyte numbers both in the lymph gland (see quantification in Supplemental Fig. 1b) and circulation (quantifications in **o**) and melanization response (quantifications in **p**). (**k, l**) Expressing *Ssadh^RNAi^* in differentiating hemocytes (*Hml^Δ^-Gal4, UAS-GFP; UAS-Ssadh^RNAi^*) leads to reduction in (**k**) circulating lamellocytes (quantifications in **q**) and (**l**) melanization response (quantifications in **r**). (**m, n**) Loss of progenitor *Gat* function (*domeMESO-Gal4, UAS-GFP; UAS-Gat^RNAi^*) leads to reduced iGABA levels both in (**m**) un-infected and (**n**) infected states. Compare with controls in **a** and **b** and see corresponding quantifications in **c.** (**o-r**) Quantifications in, (**o, p**) progenitor specific knock-down of *Gat* and *Ssadh* showing (**o**) total circulating lamellocyte numbers, *domeMESO-Gal4, UAS-GFP/+* (control, n=27), *domeMESO-Gal4, UAS-GFP; UAS-Gat^RNAi^* (n= 26, ***p<0.0001), *domeMESO-Gal4, UAS-GFP; UAS-Ssadh^RNAi^* (n= 10, ***p=0.0001) and (**p**) melanization response (Also see Table S1). (**q-r**) Differentiating cell-specific knock-down of *Gat* and *Ssadh* with (**q**) total circulating lamellocyte numbers in *Hml^Δ^-Gal4, UAS-GFP/+* (control, n=11), *Hml^Δ^-Gal4, UAS-GFP; UAS-Gat^RNAi^* (n= 17, ***p<0.0001), *Hml^Δ^-Gal4, UAS-GFP; UAS-Ssadh^RNAi^* (n= 16, ***p=0.0002) and (**r**) melanization response (Also see Table S1).

We reasoned that iGABA levels can either be dependent on GABA_B_R or a functional GABA-transporter (Gat) for its uptake(Thimgan et al., 2006) from the hemolymph. Therefore, we undertook RNA*i* based approach to manipulate expression of these respective genes and assessed for any impact on wasp-infection mediated lamellocyte differentiation and their functional role in wasp-egg melanization (See Methods for details). Independent blood-cell driver lines were employed to express RNA*i*’s directed against the respective genes either in progenitor-cells using *domeMESO-Gal4*, or in differentiating and mature blood cells of both lymph gland and circulation using *Hemolectin^Δ^-Gal4*. Following this, lamellocyte response was quantitated primarily based on their large flattened morphology identified using the cytoskeletal marker phalloidin(Small et al., 2014) and myospheroid/L4(Anderl et al., 2016). The genetically manipulated backgrounds were assessed for lymph gland responses at 24 <u>h</u>ours <u>p</u>ost infection (HPI) prior to their release into the hemolymph and 48-72 HPI for circulating blood cell counts. Multiple RNA*i* lines were tested for their knock-down efficiencies and the RNA*i* construct with the most effective knock-down efficiency was finally utilized in this study. Additional literature citing the use of these respective RNA*i* lines in other independent studies is cited in Methods section. Throughout the paper, controls refer to age-matched larvae obtained from either *w^1118^* or Gal4 genetic background crossed with *w^1118^*.

Down-regulation of *GABA_B_R* either in progenitors or differentiating cells did not affect lamellocyte development (Supplemental Fig. 1b, c). Supporting this, the formation of melanized wasp-egg capsules was detected (Supplemental Fig. 1d) and the overall cellular response to infection was comparable to control backgrounds (Supplemental Fig. 1e) as well. Thus, despite the requirement for GABA_B_R function in progenitor maintenance and steady-state lymph gland hematopoiesis(Shim et al., 2013), the immune responses seen upon infection remained unaffected by the loss of GABA_B_R signaling in blood cells. On the other hand, down-regulating *Gat* within progenitor cells (*domeMESO>Gat^RNAi^*) or differentiating blood cells (*Hml>Gat^RNAi^*) severely impaired their overall lamellocyte differentiation both in the lymph gland (Fig. 1, d, g and Supplemental Fig. 1b) and in circulation (Fig. 1, e, h and o, q). Concomitantly, the melanization response of the deposited wasp-eggs in these mutants was severely dampened as well (Fig. 1, f, i and p, r). In accordance with these genetic data, *Gat* down-regulation led to a significant reduction in blood cell iGABA levels both in homeostatic (Fig. 1c, m, Supplemental Fig. 1f) and infected conditions (Fig. 1c, n, Supplemental Fig. 1g) when compared to levels detected in controls. These data are further supported by Gat expression analysis using an anti-Gat antibody and quantitative analysis of *Gat* mRNA levels that revealed its uniform expression in lymph gland blood cells (Supplemental Fig.1h and i). The lamellocyte defect is not due to dampened cell numbers as circulating blood-cell densities detected between infected *Gat^RNAi^* mutant larvae and corresponding genetic-controls were comparable (Supplemental Fig. 1e). Thus, taken together, these data confirm a role for Gat function in blood-cell GABA uptake necessary for lamellocyte differentiation.

To elucidate if iGABA detected in blood cells is a consequence of any intracellular GABA bio-synthesis, the expression of Glutamic acid decarboxylase1 (*Gad1)*, the GABA biosynthetic enzyme was down-regulated in blood cells (*Gad1^RNAi^*). Unlike *Gat* knockdown, this genetic manipulation did not lead to any reduction in lamellocyte or melanization response (Supplemental Fig. 1c and d). On the contrary, a mild elevation in lamellocyte numbers was apparent. While this data is indicative of basal Gad1 function in blood cells but nevertheless imply a Gad1-independent but Gat-dependent control of lamellocyte differentiation.

To address the underlying cause of the lamellocyte defect in *Gat^RNAi^* background, we assayed different aspects of blood development in these animals but failed to observe any discernible changes in overall hematopoietic development or signaling implicated in blood-progenitor maintenance (Supplemental Fig. 2a-j, p, q). This analysis was undertaken by assessing overall lymph gland size, progenitor population (measured by assessing Dome-GFP^+^ area) and differentiation into mature blood cell lineages, namely plasmatocytes or crystal cells. A minor expansion of intermediate progenitors was however noticeable in *domeMESO>Gat^RNAi^* lymph glands (Supplemental Fig. 2p). Detailed expression analysis of Wingless, Hh (assessed by Cubitus interruptus expression) and Ca^2+^/GABA_B_R signaling (as assayed by CaMKII) pathways failed to reveal any changes in them (Supplemental Fig. 2c-e, h-j). Therefore, Gat function in blood cells is dispensable for steady-state hematopoiesis, but causally related to lamellocyte differentiation, the mechanistic underpinnings of which was explored next.

### iGABA metabolism via the GABA-shunt pathway controls lamellocyte differentiation

Intracellularly, GABA can be catabolized via the GABA-shunt pathway to generate succinate in two steps, which is enzymatically catalyzed by GABA-transaminase (Gabat) and Succinic-semialdehyde dehydrogenase (Ssadh)(Shelp et al., 1999). Expression analysis of *Gabat* and *Ssadh* by quantitative real-time and *in situ* hybridization based approaches detected their mRNA in blood cells (Supplemental Fig. 1i, j). As done for Gat, we manipulated the expression of these enzymes in blood cells and assessed for immune responses. Loss of Gabat in blood cells did not change lamellocyte numbers or melanization response (Supplemental Fig. 1c, d). These data were further confirmed with analyses of *Gabat ()* whole animal mutant, *gabat^PL00338^* (Maguire et al., 2015) (Supplemental Fig. 1c, d). Inhibiting the rate-limiting step of the GABA-shunt pathway catalyzed by *Ssadh*, however recapitulated the lamellocyte reduction comparable to *Gat^RNAi^* and the failure to melanize the deposited wasp-eggs (Fig. 1, j-l, o-r and Supplemental Fig. 1b). Even though lamellocyte formation was perturbed, loss of blood cell *Ssadh* function did not affect the overall circulating blood cell densities in response to wasp-infection and remained comparable to controls (Supplemental Fig. 1e). Thus, implying a specific defect in lamellocyte differentiation but not general cellular proliferation. Much like *Gat*, loss of *Ssadh* in blood cells did not alter normal hematopoiesis (Supplemental Fig. 2k-q) and implied a limiting role for Ssadh specifically for lamellocyte differentiation. The independence of *gabat* seen here in comparison to *Ssadh* cannot be explained with the data that we have, but this is not uncommon and is reported in other contexts as well(Parviz et al., 2014). In the subsequent experiments, we explored the role for Ssadh in wasp-infection mediated lamellocyte formation.

The metabolic output of Ssadh enzymatic reaction is the generation of succinate (Fig. 2a). We explored if the lack of succinate in the GABA-shunt pathway mutants was responsible for the lamellocyte defect and if supplementing succinate to *Drosophila* larvae expressing *Gat^RNAi^* or *Ssadh^RNAi^* in both *domeMESO>* and *Hml^Δ^>* genetic backgrounds, corrected their lamellocyte defects. For this, first instar larvae were raised on food containing 3-5% succinate and then subjected to wasp-infections followed by analysis of their cellular immune response. Compared to mutants raised in regular food, the succinate supplemented diet significantly restored circulating and lymph gland lamellocyte numbers (Fig. 2, b-e, h, i and Supplemental Fig. 3a). Consequently, lamellocyte enwrapped wasp-eggs (Supplemental Fig. 3, b-e) and the formation of melanotic capsules were evident (Fig. 2, f, g, j, k) in the rescued animals. Succinate supplementation did not affect overall larval health nor did it alter blood development as assessed by characterization of lymph gland hematopoietic profile (Supplemental Fig. 3f, g). The rescue is specifically seen with succinate and not detected with other similar metabolites like α-ketoglutarate (α-KG) and fumarate (Supplemental Fig. 3h). Together, these data reveal a role for GABA-shunt pathway in supporting succinate availability in blood cells that is necessary to mount a cellular lamellocyte response upon wasp-infections.

**Figure 2.**
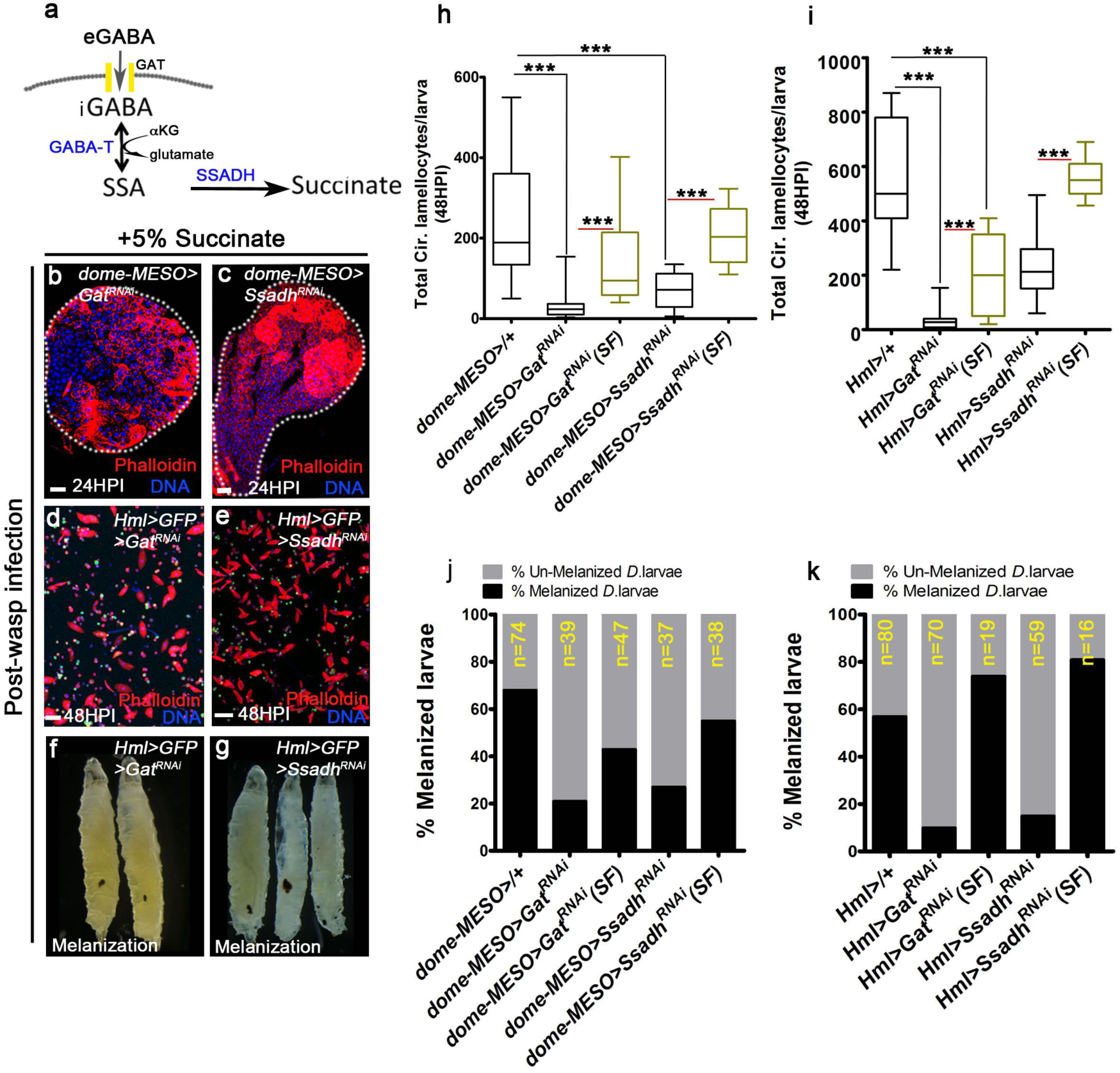
GABA-shunt derived succinate controls lamellocyte potential. DNA is marked with DAPI (blue) and blood cells are marked with phalloidin (red). In panels **b, c** scale bars = 20μm and **d, e** = 50μm. HPI indicates <u>h</u>ours <u>p</u>ost wasp-infection, RF is regular food and SF is succinate food. Statistical analysis in **h** and **i** is Mann-Whitney test, two-tailed. In panels **h, i** median is shown in box plots and vertical bars represent upper- and lowest cell-counts. **“**n” is total number of animals analyzed. **(a)** A schematic of the GABA-shunt pathway. Uptake of extra-cellular GABA (eGABA) via Gat (yellow bars) in blood cells and its intracellular catabolism through GABA-transaminase (GABA-T) which catalyzes the conversion of GABA into succinic semi-aldehyde (SSA) and its further breakdown into succinate by succinic semi-aldehyde dehydrogenase (SSADH). **(b-g)** 5% succinate supplementation restores (**b-e**) lamellocyte and (**f, g**) melanization defect of (**b, d** and **f)** *Gat^RNAi^* or (**c, e** and **g**) *Ssadh^RNAi^* mutants. See quantifications of the respective genotypes in **h-k**. **(h, i)** Total circulating lamellocyte quantifications in (**h**) *domeMESO-Gal4, UAS-GFP/+* on RF (control, n=27), *domeMESO-Gal4, UAS-GFP; UAS-Gat^RNAi^* on RF (n= 26, ***p<0.0001), *domeMESO-Gal4, UAS-GFP; UAS-Gat^RNAi^* on SF (n= 12, ***p<0.0001), *domeMESO-Gal4, UAS-GFP; UAS-Ssadh^RNAi^* on RF (n= 10, ***p=0.0001) and *domeMESO-Gal4, UAS-GFP; UAS-Ssadh^RNAi^* on SF (n= 6, ***p=0.001). (**i**) *Hml^Δ^-Gal4, UAS-GFP/+* on RF (control, n=11), *Hml^Δ^-Gal4, UAS-GFP; UAS-Gat^RNAi^* on RF (n= 17, ***p< 0.0001), *Hml^Δ^-Gal4, UAS-GFP; UAS-Gat^RNAi^* on SF (n=11, ***p=0.001), *Hml^Δ^-Gal4, UAS-GFP; UAS-Ssadh^RNAi^* on RF (n= 16, ***p=0.0002) and *Hml^Δ^-Gal4, UAS-GFP; UAS-Ssadh ^RNAi^* on SF (n= 7, ***p=0.0003). **j, k**) Graphical representation of melanization response in *Gat^RNAi^* and *Ssadh^RNAi^* animals following succinate supplementation (refer Table S1).

### TCA-independent control of lamellocyte response

Succinate is also derived from the tricarboxylic acid cycle (TCA) via the conversion of α-ketoglutarate which is catalyzed in a two-step process by *α-ketoglutarate dehydrogenase, αKDH* (*CG33791*)(Zhou et al., 2008) and *succinyl CoA synthetase, skap (CG11963)*(Gao et al., 2008). Unlike modulation of the GABA-shunt pathway, down-regulating these TCA enzymes did not lead to any defect in lamellocyte formation or melanization of the deposited wasp-eggs upon wasp-infection (Supplemental Fig. 4a-e). Even though, expression analysis through quantitative and *in situ* based approaches detected their expression in control lymph glands (Supplemental Fig. 4f, g), loss of *αKDH* or *skap* function in blood cells did not reveal any changes in overall blood development either (Supplemental Fig. 4h-m). Therefore, suggestive of a TCA-independent but GABA-shunt dependent control of lamellocyte differentiation and normal blood development. The independence of TCA in this context is intriguing and we speculate separate pools of succinate in blood cells that are maintained to control basal cellular metabolism and specialized immune requirements. The TCA-derived succinate most-likely conducts basal metabolic functions and the GABA-shunt derived succinate sustains the immune requirement of these blood cells. As a result, blocking GABA-shunt pathway without compromising basal cellular metabolism still allows the development of blood cells, but nevertheless impedes lamellocyte potential.

Next, the downstream effects of succinate responsible for lamellocyte differentiation was investigated. As a metabolite, succinate fuels the activity of Succinate dehydrogenase (SDH) complex, which is the complex II of the mitochondrial respiratory chain that converts succinate to fumarate(Rutter et al., 2010). Inhibiting SDH function by blocking its catalytic sub-unit SdhA (*SdhA^RNAi^*) as the means to prevent succinate utilization within the TCA failed to cause any reduction in lamellocyte differentiation or melanization response. On the contrary a stark increase was detected (Supplemental Fig. 5a-c), which we hypothesize could be an outcome of further elevation in succinate levels in these animals. Thereby implying, an alternative role for succinate in lamellocyte differentiation which is enhanced in *SdhA^RNAi^* background.

### Progenitor Sima protein stability via GABA-shunt establishes lamellocyte immune-potential

Multiple studies across model systems and cell types have reported an integral role for succinate in hypoxia-independent stabilization of Hypoxia-inducible factor (HIFα) via inhibition of prolyl hydroxylases that mark HIFα protein for degradation(Briere et al., 2005; Selak et al., 2005; Tannahill et al., 2013). Within the *Drosophila* larval hematopoietic tissue, Sima protein, orthologous to mammalian HIFα(Lavista-Llanos et al., 2002) is detected at basal levels in all cells of the larval lymph gland with comparatively higher expression in crystal cells as previously reported(Mukherjee et al., 2011) (Fig. 3a). Within a few hours of wasp-infection (6HPI) the basal expression of Sima protein in lymph gland blood cells is up-regulated (Fig. 3b, c, Supplemental Fig. 5d). This is prior to detection of any lamellocyte formation. Later, as lamellocyte formation ensues (12HPI onwards), Sima protein expression is detected in them as well (Supplemental Fig. 5e). A similar increase in sima *mRNA* levels is however not apparent (Supplemental Fig. 5f), indicating either a translational or post-translational control of Sima protein expression in blood cells upon wasp-infections. Modulating blood-progenitor and differentiating blood cell-specific expression of Sima by employing *sima^RNAi^* severely impaired lamellocyte formation (Fig. 3d and Supplemental Fig. 5g) and wasp-egg melanization response (Supplemental Fig. 5h). Gain of *sima* function on the other hand caused an elevation in lamellocyte production (Supplemental Fig. 5g, h). Thereby, indicating a necessary and sufficient role for Sima in lamellocyte induction. As seen previously for GABA shunt pathway components, loss of Sima function also did not reveal any significant defect in lymph gland progenitor homeostasis or their differentiation status in normal uninfected conditions but an increase in intermediate progenitor population was evident (Supplemental Fig. 5i-k). Genetic perturbations to manipulate Sima protein stability via regulating hydroxy-prolyl hydroxylase (Hph(Schofield and Ratcliffe, 2005)) expression also recapitulated the lamellocyte defects. Hph expression in blood-progenitor or differentiating cells was either elevated (*UAS-Hph*) or down-regulated (*Hph^RNAi^*). These led to corresponding loss or gain of lamellocyte numbers respectively (Fig. 3d and Supplemental Fig. 5, g, h), implying an important role for immune-cell Hph activity in moderating lamellocyte response.

**Figure 3.**
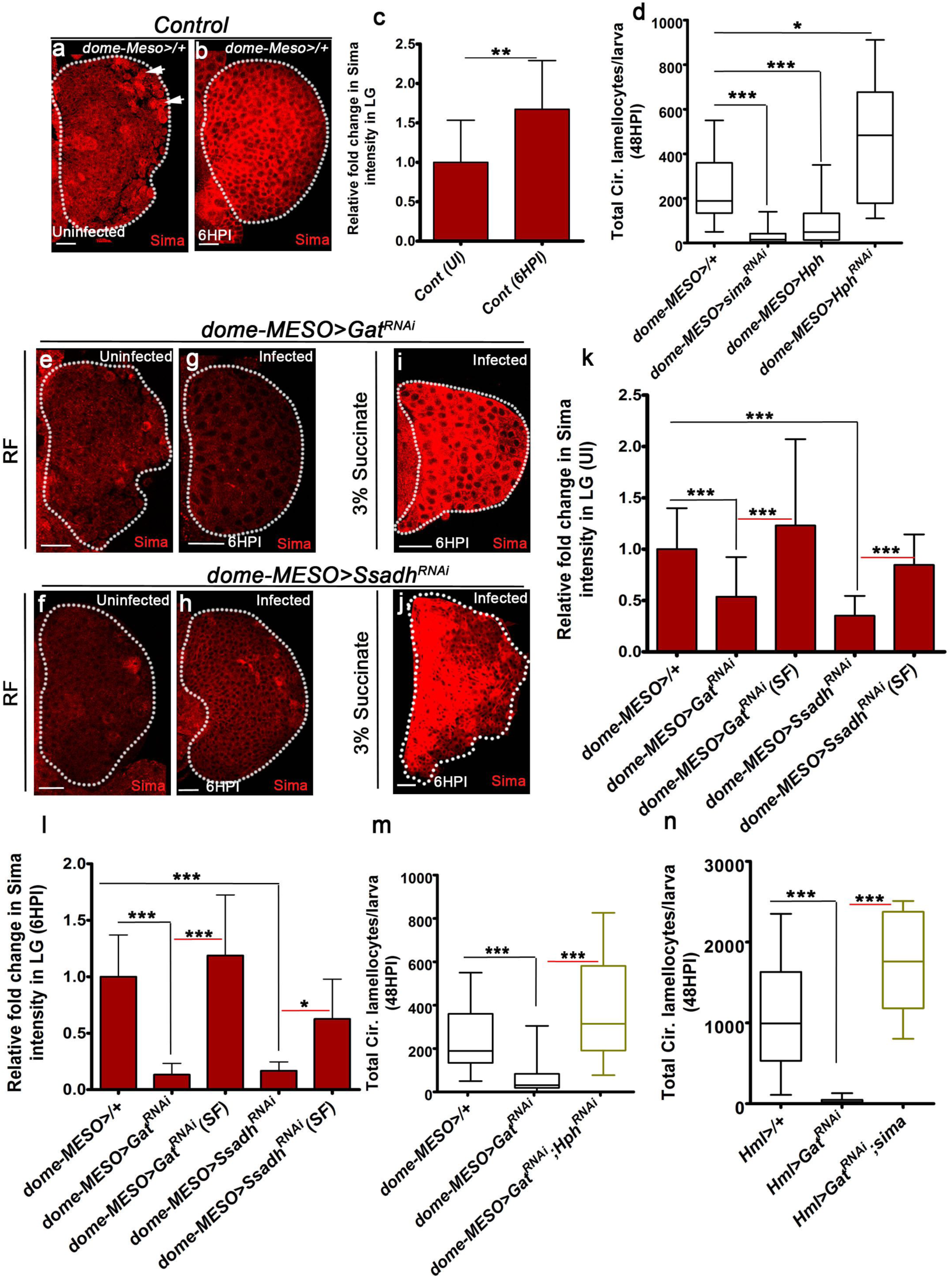
GABA-shunt dependent control of Sima protein stabilization in immune-progenitor cells promotes lamellocyte induction. In all panels Sima protein is in red. Scale bars = 20μm. UI is uninfected, HPI is <u>h</u>ours <u>p</u>ost wasp-infection, RF is regular food and SF is succinate food. Bars in panel **c, k, l** represent standard error of mean (SEM). In **d, m, n** median is shown in box plots and vertical bars represent upper and lowest cell-counts. Statistical analysis, in panels **c, k, l** is un-paired *t*-test, two-tailed and in panels **d, m, n**, Mann-Whitney test, two-tailed. “n” represents total number of larvae analyzed. (**a-c**) Compared to Sima expression detected in control (*domeMESO>GFP/+*) lymph glands obtained from (**a**) un-infected animals (crystal cells marked with white arrows), **(b)** Sima is elevated at 6HPI. (**c**) Quantification of relative fold change in Sima intensities in UI (n=18), 6HPI (n=18, **p=0.0013). See Supplemental Fig. **5d** for mean Sima intensities. (**d**) Quantitative representation of circulating lamellocyte counts in *domeMESO-Gal4, UAS-GFP*/+ (control, n=27), *domeMESO-Gal4, UAS-GFP*; *UAS-sima^RNAi^* (n=29, ***p<0.0001), *DomeMESO-Gal4, UAS-GFP*; *UAS-Hph* (n=22, ***p<0.0001) and *DomeMESO-Gal4, UAS-GFP*; *UAS-Hph^RNAi^* (n=18, *p=0.01). (**e, f**) Compared to Sima levels detected in developing un-infected lymph glands from (**a)** control larvae, **(e)** *domeMESO-Gal4, UAS-GFP*; *UAS-Gat^RNAi^* and (**f**) *domeMESO-Gal4, UAS-GFP*; *UAS-Ssadh^RNAi^* show reduction in Sima expression. Quantified in **k** and Supplemental Fig. 6**a, a’**. (**g-j**) Compared to Sima levels seen post-infection in lymph glands on RF from (**b)** control, **(g)** *domeMESO-Gal4, UAS-GFP*; *UAS-Gat^RNAi^* and (**h**) *domeMESO-Gal4, UAS-GFP*; *UAS-Ssadh^RNAi^* show reduction that is (**i, j**) restored with succinate (SF). (**i)** *domeMESO-Gal4, UAS-GFP*; *UAS-Gat^RNAi^* on SF, (**j)** *domeMESO-Gal4, UAS-GFP*; *UAS-Ssadh^RNAi^* on SF. Quantified in **l** and Supplemental Fig. **6b**. (**k, l**) Quantifications of relative fold change in lymph gland Sima intensities in (**k**) un-infected and (**l**) infected 6HPI conditions. (**k**) *DomeMESO>GFP/+* on RF (control, n=25), *domeMESO-Gal4, UAS-GFP/UAS-Gat^RNAi^* on RF (n=29, ***p<0.0001) and *domeMESO-Gal4, UAS-GFP/UAS-Gat^RNAi^* on SF (n= 16, ***p=0.0003), *domeMESO-Gal4, UAS-GFP/UAS-Ssadh^RNAi^* on RF (n=18, ***p<0.0001) and *domeMESO-Gal4, UAS-GFP/UAS-Ssadh^RNAi^* on SF (n= 15, ***p<0.0001). 6HPI (**l**) *domeMESO-Gal4>/+* on RF (control, n=18), *domeMESO-Gal4, UAS-GFP/UAS-Gat^RNAi^* on RF (n=18, ***p<0.0001) and *domeMESO-Gal4, UAS-GFP/UAS-Gat^RNAi^* on SF (n= 9, ***p<0.0001), *domeMESO-Gal4, UAS-GFP/UAS-Ssadh^RNAi^* on RF (n=6, ***p<0.0001) and *domeMESO-Gal4, UAS-GFP/UAS-Ssadh^RNAi^* on SF (n= 7, *p=0.01). See Supplemental Fig.**6 a-b** for mean Sima intensities. (**m, n**) Quantification of total circulating lamellocyte counts in (**m**) *domeMESO-Gal4>/+* (control, n=27), *domeMESO-Gal4, UAS-GFP/UAS-Gat^RNAi^* (n=24, ***p<0.0001) and *domeMESO-Gal4, UAS-GFP/UAS-Gat^RNAi^; UAS-Hph^RNAi^* (n= 9, ***p=0.0001) and (**n**) *Hml^Δ^-Gal4, UAS-GFP/+* (control, n=10), *Hml^Δ^-Gal4, UAS-GFP; UAS-Gat^RNAi^* (n=16, ***p<0.0001) and *Hml^Δ^-Gal4, UAS-GFP/UAS-sima; UAS-Gat^RNAi^* (n= 10, ***p<0.0001).

Based on all the above-mentioned genetic and Sima protein expression data, we hypothesized a role for GABA-shunt in Sima protein stabilization via succinate mediated inhibition of Hph activity. We tested this hypothesis by first employing tools to examine Sima protein expression in GABA-shunt mutants followed by genetic epistasis tests. Sima protein in *Gat^RNAi^* and *Ssadh^RNAi^* mutant lymph glands was analyzed during development and post wasp-infection. Compared to 3^rd^ instar control lymph glands obtained from uninfected larvae, Sima protein detected in *domeMESO>Gat^RNAi^* and *domeMESO>Ssadh^RNAi^* mutants was significantly low (Fig. 3, e, f, k and Supplemental Fig. 6, a, a’). These mutants also demonstrated a failure to raise Sima protein post-infection (Fig. 3, g, h and l, Supplemental Fig. 6b). Succinate supplementation of *Gat^RNAi^* and *Ssadh^RNAi^* mutants significantly restored Sima protein levels comparable to controls both in uninfected and infected states (Fig. 3, i-l and Supplemental Fig. 6, a-b). To validate if succinate mediated control of Sima protein expression was indeed at the level of inhibiting Hph activity, *Hph^RNAi^* was expressed in blood progenitor-cells lacking *Gat* expression (*domeMESO>UAS-Gat^RNAi^*; *UAS-Hph^RNAi^*). This genetic combination restored lamellocyte numbers almost comparable to infected controls (Fig. 3m). Direct over-expression of *sima* in differentiating blood cells expressing *Gat^RNAi^* (*Hml>UAS-sima; UAS-Gat^RNAi^*) also restored their lamellocyte numbers (Fig. 3n). Notably, the lamellocytes generated in the aforementioned genetic experiments displayed wasp-egg encapsulation (Supplemental Fig. 6c), however the melanization response in these animals was undetectable Supplemental Fig. 6d). Post-infection, succinate supplementation of *Hml>Gat^RNAi^*; *UAS-sima* corrected the melanization defect normally undetectable in *Hml>Gat^RNAi^* only backgrounds (Supplemental Fig. 6d). Thus, alluding to a Sima independent role for succinate in melanization response as well.

Down-stream of Sima we investigated the canonical hypoxia response pathway which is mediated by interaction with tango (tgo)/β-ARNT partner necessary for hypoxia response(Schofield and Ratcliffe, 2005). Expressing *tgo^RNAi^* in the blood cells failed to cause any defect in lamellocyte differentiation or melanization response (Supplemental Fig. 6, e-h). Independent of Sima interaction with tgo, previous findings have also reported non-canonical interaction with Notch necessary for blood cell survival(Mukherjee et al., 2011). Loss of blood-cell Notch function in progenitor or differentiating cells also did not recapitulate any lamellocyte or melanization defect (Supplemental Fig. 6, e-h). When assessed for the requirement of a downstream target gene controlled by Sima, namely *lactate dehydrogenase (Ldh)*(Lavista-Llanos et al., 2002) we observed a strong requirement for its function in lamellocyte and melanization response (Supplemental Fig. 6, e-h). Consistent with its functional requirement, lymph gland mRNA samples obtained from infected larvae revealed a 30-fold increase in *Ldh mRNA* expression (Supplemental Fig. 6i) as also reported previously(Bajgar et al., 2015). Together, these findings confirm a requirement for Ldh in wasp-infection mediated lamellocyte response, but the mechanism by which Sima functions independent of Tgo and Notch needs to be investigated further.

Overall, the data thus far highlight an immune cell mediated uptake of systemic GABA in the development of competent blood-progenitor cells. iGABA catabolism via the GABA-shunt pathway generates succinate, which is necessary to mediate two functions. Succinate during hematopoiesis controls Hph activity in blood-progenitors and differentiating cells which is necessary for the stabilization and maintenance of basal Sima protein in them. Sima establishes basic competency of immune cells to generate lamellocytes. Following infections, iGABA levels rise and this leads to further elevation of Sima protein to most-likely achieve a critical threshold that triggers lamellocyte induction. Secondary to Sima stabilization, the succinate derived from GABA is also required for activating the melanization cascade. Together, this ascertains a successful immune response.

The critical control of systemic GABA uptake by immune cells for wasp-mediated immune response warrants an understanding of upstream events mediating GABA availability. Our past work has implicated olfactory inputs in controlling system-level changes of GABA. This is primarily the GABA that is detected by the immune-progenitor cells. We explored the control of olfactory inputs in cellular immune response to wasp-infection.

### Olfaction-derived GABA controls blood-progenitor immune-competency

During larval development, we know that sensing food-odors by specific larval olfactory receptor neurons (ORNs) leads to downstream activation of projection neurons (PNs). This stimulates a subset of neurosecretory cells (Kurs6^+^) in the central brain to synthesize and secrete GABA into the hemolymph which is sensed by the blood progenitor cells(Shim et al., 2013). We found that abrogating olfaction mimicked the lamellocyte differentiation phenotype and wasp-egg melanization defects (Fig. 4, a-i and Supplemental Fig. 7, a-d). This was genetically addressed by: 1) analyzing *orco* mutant(Neuhaus et al., 2005), the common odorant co-receptor 83b necessary for all odor responsiveness(Larsson et al., 2004) (*orco^1^/orco^1^*; Fig. 4, a-e and Supplemental Fig. 7, l, m), (2) ablating all ORNs (*Orco>Hid, rpr* Supplemental Fig. 7, a, b) and (3) specific ablation of Or42a, the ORN implicated in sensing food related odors (*OR42a>Hid*, Fig. 4, f-i). Physiological experimental set up designed to test the involvement of odors also led to the same conclusion and phenocopied lamellocyte and melanization defects (Supplemental Fig. 7, c, d). This was addressed by rearing *Drosophila* larvae from an early embryonic stage in food-medium with minimal food odors but nutritionally equivalent to regular food. Restoring food odors back into the minimal odor medium restored the cellular immune defects significantly (Supplemental Fig. 7, c-d). Downstream of odor-sensing, blocking projection neurons by inhibiting their acetylcholine synthesis (*GH146>ChAT^RNAi^*, Supplemental Fig. 7, e, f) or blocking GABA biosynthesis in Kurs6^+^ neurosecretory cells (*Kurs6>Gad1^RNAi^*, Fig. 4, j-l) phenocopied the lamellocyte defect. As a result a strong reduction in melanization response was noted in *Kurs6>Gad1^RNAi^* animals (Fig. 4, j” and m) and a comparatively milder reduction was seen in *GH146>ChAT^RNAi^* (Supplemental Fig. 7f). Like *Gat* and *Ssadh*, none of these above-mentioned neuronal conditions impeded differentiation of other blood-cell lineages, nor did they affect overall circulating blood cell numbers following infections (Supplemental Fig. 7, g-k). Thus, based on all these data it is apparent that odor-sensing via Or42a mediates stimulation of neuronal GABA production which is sufficient for priming lamellocyte potential of blood progenitor cells. These data however do not rule out the involvement of other ORs functioning synergistically to control cellular immune response. GABA or succinate supplemented diet significantly restored the lamellocyte defects detected in animals with either olfactory dysfunction (*orco^1^/orco^1^, Orco>Hid, rpr*, Fig. 5, a, b and Supplemental Fig. 8a) or animals with neuronal loss of GABA biosynthesis (*Kurs6>Gad1^RNAi^*, Fig. 5, c, d). Representative images in Fig. 5, e-h’ highlight the restoration of lamellocytes in the lymph glands and circulation of the mutants with GABA (Fig. 5, e, e’, g, g’) or succinate (Fig. 5, f, f’, h, h’) supplementation respectively. Consistent with the requirement of the neuronal inputs in lamellocyte formation, a significant reduction in Sima protein expression was apparent in blood cells from the different neuronal mutant conditions (Fig. 6, b, d compared to control in a and Supplemental Fig. 8, b-e) which was restored in a succinate supplemented diet (Fig. 6, a-f and Supplemental Fig. 8, c-f). Further corroborating with these data, the lamellocyte defect seen in *orco* mutant or in minimal odor food was reverted by either blocking *Hph* expression in blood cells (Fig. 6, g-j) or by restoring blood cell Sima protein expression (Supplemental Fig. 8g). Figure 6g shows a representative lymph gland image depicting differentiating blood cells obtained from *orco* mutant larvae and their failure to generate lamellocytes (in red). In these animals blood-cell specific expression of *Hph^RNAi^* rescued the lamellocyte defect significantly (Fig. 6h). Thus, proving a critical dependence of the larval hematopoietic system on olfactory stimulation for GABA production necessary to prime immune-cell potential to generate lamellocytes. This systemic metabolic axis functions in addition to GABA/GABA_B_R dependent control of blood-progenitor development and maintenance that is previously reported(Shim et al., 2013).

**Figure 4.**
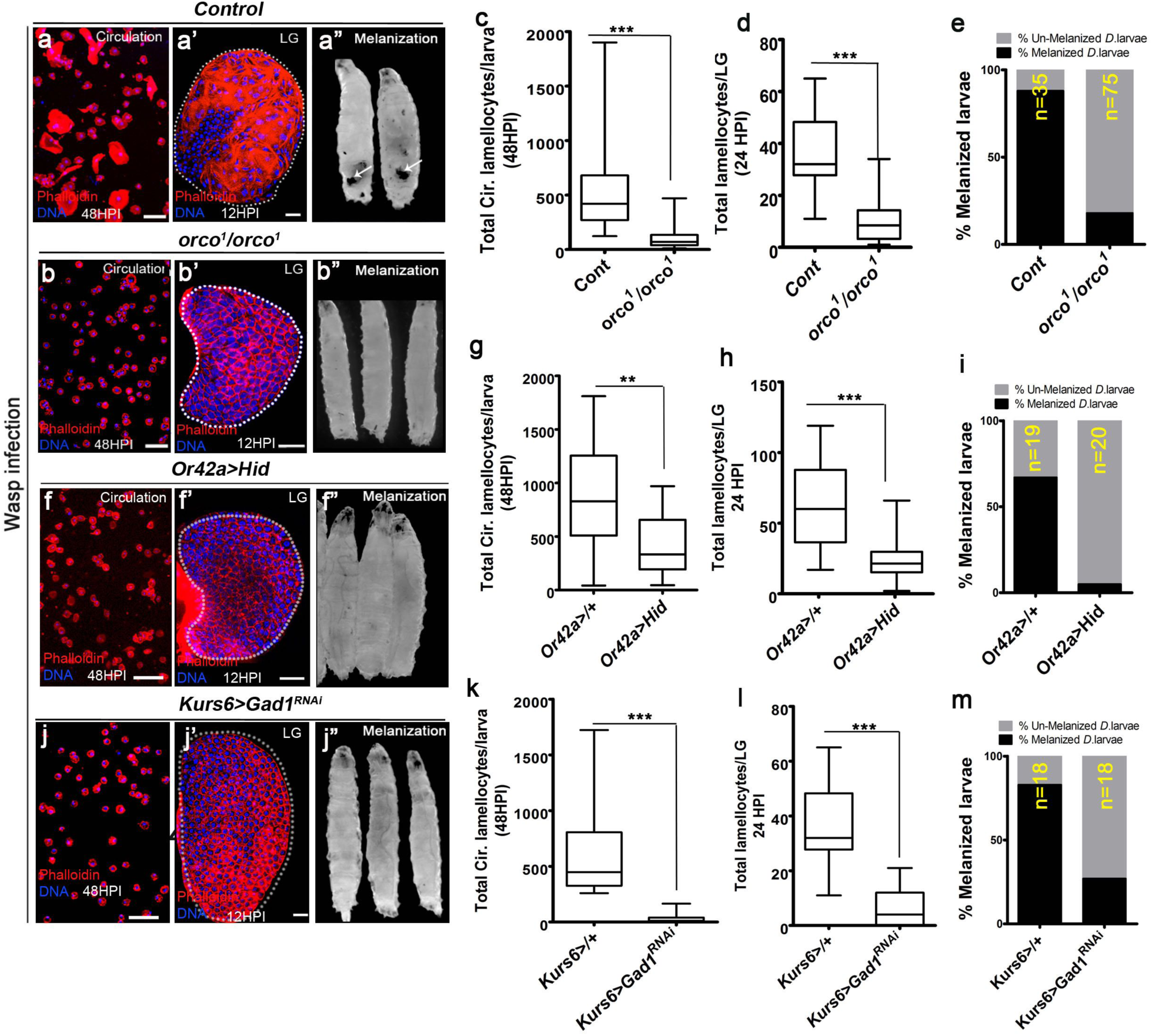
Odor-mediated neuronal GABA availability specifies lamellocyte potential. DNA is stained with DAPI (blue). Phalloidin (red) marks blood cells and lamellocytes are characterzied by their large flattened morphology. Scale bars in panels **a**, **b, f, j** = 50μm and **a’, b’, f’, j’** = 20μm. HPI indicates <u>h</u>ours <u>p</u>ost wasp-infection, RF is regular food. In panels **c, d, g**, **h**, **k, l** median is shown in box plots and vertical bars represent upper and lowest cell-counts. Statistical analysis, in panels **c, d, g**, **h**, **k, l** Mann-Whitney test, two-tailed. “n” represents the total number of larvae analyzed. **(a-a’’)** Control (*w^1118^*) infected larvae showing lamellocyte induction in (**a**) circulation, (**a’**) lymph gland and (**a”**) melanization. Quantifications in **c-e**. **(b-e)** Compared to (**a-a”**) control (*w^1118^*), *orco^1^/orco^1^* mutant larvae show reduction in lamellocyte in (**b, c**) circulation (n=18, ***p<0.0001 compared to *w^1118^*, n= 23) and (**b’, d**) lymph glands (n=20, ***p<0.0001 compared to *w^1118^*, n= 18) along with (**b”, e**) reduced melanization. **(f-i)** Specifically ablating Or42a *(Or42a-Gal4, UAS-Hid)* causes reduction in lamellocytes in (**f, g**) circulation (n=20, ** p= 0.004 compared to *Or42a-Gal4*/+, n=19) and (**f’, h**) lymph gland (n=24, ***p<0.0001 compared to *Or42a-Gal4*/+, n= 24) and (**f’’, i**) along with reduced melanization. **(j-m)** Blocking neuronal GABA bio-synthesis in Kurs6^+^ neurons (*Kurs6-Gal4; UAS-Gad1^RNAi^*) recapitulates lamellocyte reduction in (**j, k**) circulation (n=25, ***p<0.0001 compared to *Kurs6-Gal4*/+, n= 22), (**j’, l**) lymph gland (n=22, ***p<0.0001 compared to *Kurs6-Gal4*/+, n= 18) and (**j”, m**) reduced melanization.

**Figure 5.**
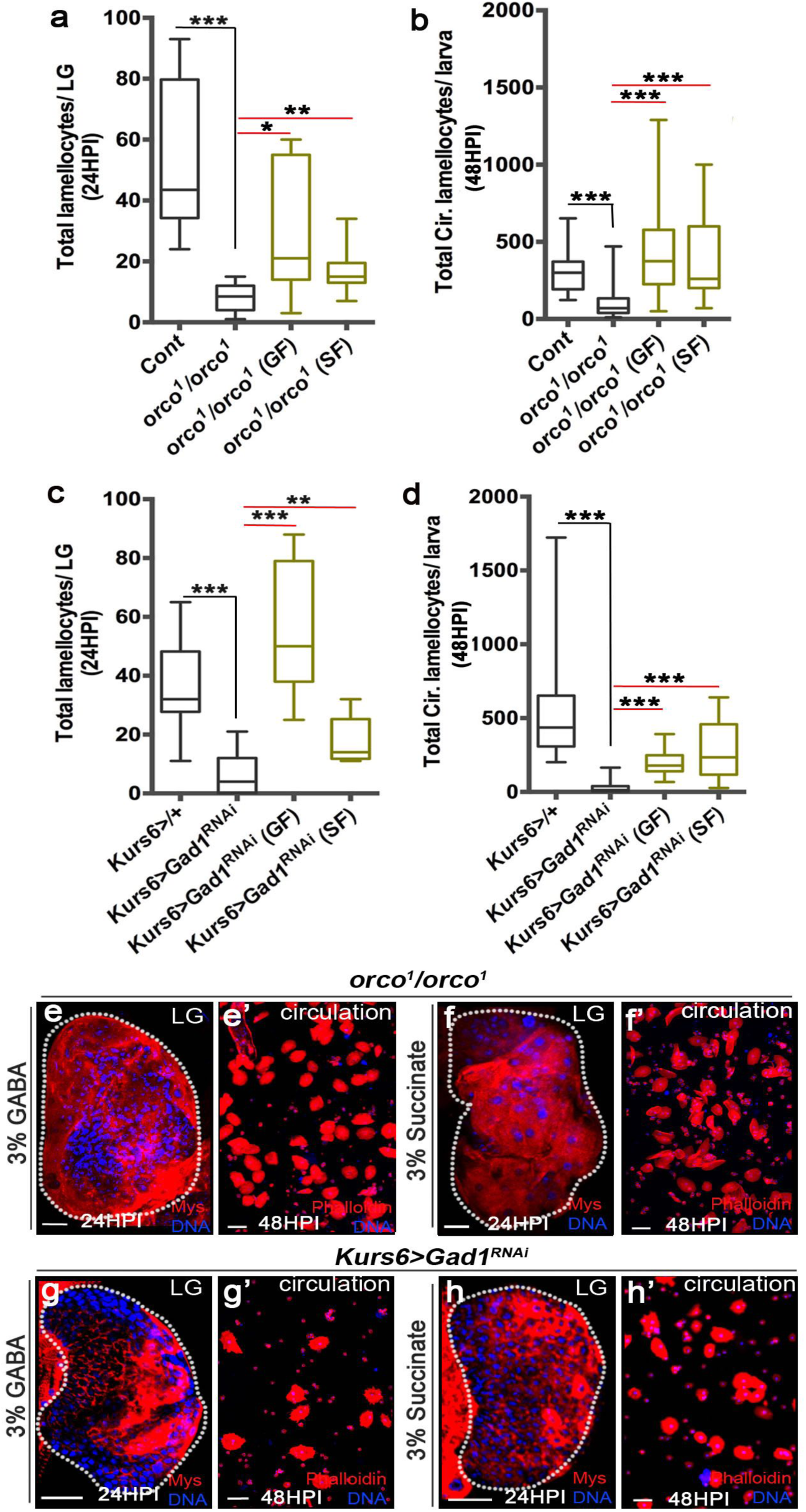
Olfaction derived GABA controls blood cell succinate levels. DNA is stained with DAPI (blue). Lamellocytes in panels **e, f, g, h** are marked with Myospheroid (red) and **e’, f’, g’, h’** with Phalloidin (red). Lamellocytes are characterzied by their large flattened morphology. Scale bars in panels **e**, **f, g, h** = 20μm and **e’, f’, g’, h’** = 50μm. HPI indicates <u>h</u>ours <u>p</u>ost wasp-infection, RF is regular food, GF is GABA supplemented food, SF is succinate supplemented food. Mann-Whitney test, two-tailed has been applied for statistical analysis. Median plotted in box plots and vertical bars represent upper and lowest cell-counts. “n” represents the total number of larvae analyzed. (**a-d**) Quantification of total lamellocyte counts in (**a, c**) lymph glands and (**b, d**) circulation. (**a**) *orco^1^/orco^1^* mutant in RF (n=10, ***p<0.0001 compared to control, *w^1118^*, n=8), GF (n= 7, *p=0.02), SF (n=13,**p=0.006). (**b**) *orco^1^/orco^1^* mutant in RF (n=41, ***p<0.0001 compared to control, w^1118^, n=14), GF (n= 18, ***p<0.0001), SF (n=14, ***p<0.0001). (**c**) *Kurs6-Gal4, UAS-Gad1^RNAi^* in RF (n=22, ***p<0.0001 compared to *Kurs6-Gal4/+*, control n=18), GF (n=11 and ***p<0.0001) and SF (n=6, **p=0.007). (**d**) *Kurs6-Gal4, UAS-Gad1^RNAi^* in RF (n=25, ***p<0.0001 compared to *Kurs6-Gal4/+*, control n=25), GF (n=12 and ***p<0.0001) and SF (n=26, ***p<0.0001). (**e-f’**) Representative images of lamellocyte rescue with (**e, e’, g, g’**) GABA and (**f, f’, h, h’**) succinate in lymph gland and circulation of (**e-f’**) *orco^1^/orco^1^* and (**g-h’**) *Kurs6-Gal4, UAS-Gad1^RNAi^* larvae.

**Figure 6.**
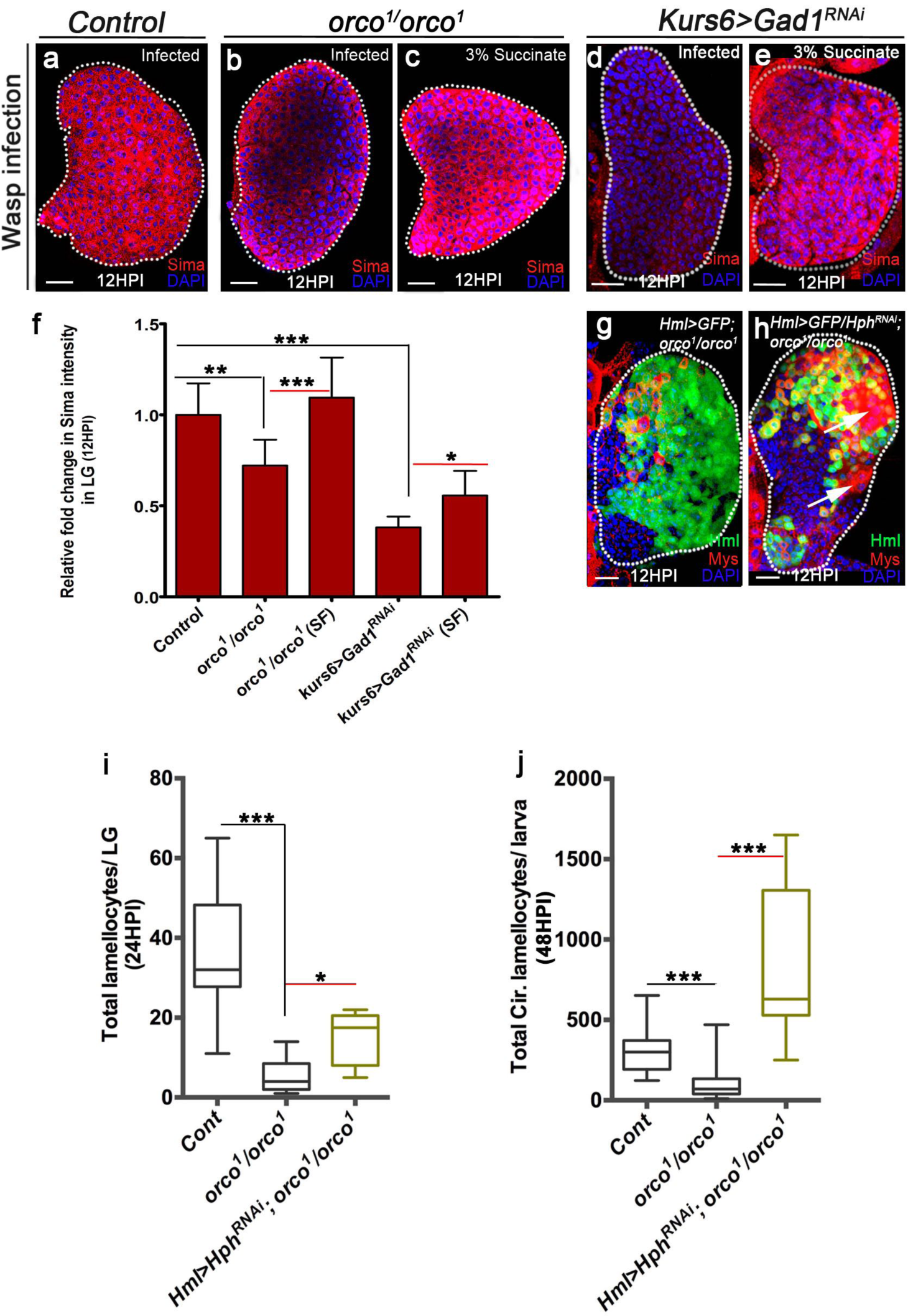
Olfaction controls Sima protein expression in blood cells. DNA is stained with DAPI (blue). Shown in panels **a-e** Sima protein (red). In **g, h** Myospheroid (red) marks lamellocytes which are characterzied by their large flattened morphology. Scale bars = 20μm. HPI indicates <u>h</u>ours <u>p</u>ost wasp-infection, RF is regular food SF is succinate supplemented food. Panel **f** represents fold change with standard error of mean (SEM). In **i, j** median is shown in box plots and bars represent upper and lowest cell-counts. Statistical analysis, in panel **f** un-paired *t*-test, two-tailed and in panels **i, j** Mann-Whitney test, two-tailed. “n” represents total number of larvae analyzed. (**a-f**) Compared to Sima protein levels detected at 12HPI in lymph glands on RF from (**a**) control animals (*w^1118^*), (**b**) *orco^1^/orco^1^* and (**d**) *Kurs6-Gal4, UAS-Gad1^RNAi^* show reduced Sima expression which is (**c, e**) restored on SF, (**c**) *orco^1^/orco^1^* on SF and (**e**) *Kurs6-Gal4, UAS-Gad1^RNAi^* on SF. (**f**) Corresponding quantifications of the relative fold change in Sima intensities in these backgrounds. Control on RF (*w^1118^*, n=9), *orco^1^/orco^1^* on RF (n=9, **p=0.002) and *orco^1^/orco^1^* on SF (n= 10, ***p=0.0004), *Kurs6-Gal4, UAS-Gad1^RNAi^* on RF (n=6, ***p<0.0001) and *Kurs6-Gal4, UAS-Gad1^RNAi^* on SF (n= 8, *p=0.01). See Supplemental Fig. **8c** for mean intensity values. (**g-j**) Blood cell specific loss of *Hph* function in *orco* mutants restores lamellocyte response both in the (**g-i**) lymph gland and (**j**) circulation. Representative lymph gland images of (**g**) *Hml^Δ^-Gal4, UAS-GFP; orco^1^/orco^1^* lacking lamellocytes (as evident by lack of large flattened cells) but contain differentiating immune cells (Hemolectin, green). (**h**) In *Hml^Δ^-Gal4, UAS-GFP, UAS-Hph^RNAi^*; *orco^1^/orco^1^* genetic background, lamellocyte population (marked with white arrows) are detected. (**i, j**) Quantification of lamellocytes in (**i**) lymph glands, control (*w^1118^*, n=18), *orco^1^/orco^1^* (n=10, ***p<0.0001), *Hml>Hph^RNAi^; orco^1^/orco^1^* (n= 6, *p=0.01) and (**j**) circulation, control (*w^1118^*, n=14), *orco^1^/orco^1^* (n=18, ***p<0.0001) and *Hml^Δ^-Gal4, UAS-GFP, UAS-Hph^RNAi^*; *orco^1^/orco^1^* (n=11, ***p<0.0001).

### Pathogenic odors influence blood immune potential

The establishment of lamellocyte potential by a long-range metabolic cross-talk set up by odor detection is rather puzzling. We asked if the olfactory axis was involved in sensing wasps. We addressed this by employing a physiological approach. For this *Drosophila* larvae were reared from early embryonic stages in a food medium that was infused with wasp odors (condition referred to as WOF and see Methods). This was undertaken to mimic larvae rearing in wasp-infested scenarios as would be expected in the wild, where the chances of infection are higher. If prior pathogenic odor experience during development influenced any aspect of blood development and immune response was examined. The pre-conditioned animals were subjected to wasp immune challenge with *L.boulardi* (see Methods) followed by analysis of their cellular immune response. For comparisons, immune response of infected larvae reared in regular food medium was used as experimental controls. We found that WOF animals demonstrated a significant increase in lamellocyte numbers (Fig. 7a and Supplemental Fig. 9a). Almost 1.5 to a 2 fold increase in lamellocyte numbers was evident in wasp-infected larvae raised in wasp-odor enriched condition (Fig. 7a and Supplemental Fig. 9a). A significant increase in hemolymph GABA levels was also noticable (Fig. 7b and Supplemental Fig. 9b). While the effects observed are mild, the increase in lamellocyte and GABA levels are consistently detected in distinct genetic backgrounds (Fig. 7, a, b and Supplemental Fig. 9, a, b) and completely diminished in *orco* mutant animals (Supplemental Fig. 9a). Implying an olfaction dependent control of wasp odor-detection in elevating immune response that is most likely not a consequence of contact or ingestion of wasp-odor components. Moreover, the lamellocyte sizes detected at 12HPI in lymph glands from animals reared in WOF pre-conditioned food in comparison to animals grown on regular food medium, were much larger (Fig. 7d compared to control media in c). The WOF animals also demonstrated improved melanization efficiencies (Supplemental Fig. 9c, see Methods). A mild to moderate increase in lymph gland blood-cell iGABA (Fig. 7, f compared to e, l and Supplemental Fig. 10a and Supplemental Fig. 9d) and Sima protein (Fig. 7h compared to g, m and Supplemental Fig. 10a and Supplemental Fig. 9d) expression was also detected. Thus, combining all the approaches employed that include: lamellocyte numbers, melanization response, hemolymph GABA levels, lymph gland measure of GABA and Sima levels, we observed that pre-conditioning *Drosophila* larvae with wasp odors impacts larval hematopoiesis. The wasp-odors enhanced immune-competency of blood progenitor cells and these animals generated lamellocytes efficiently.

**Figure 7.**
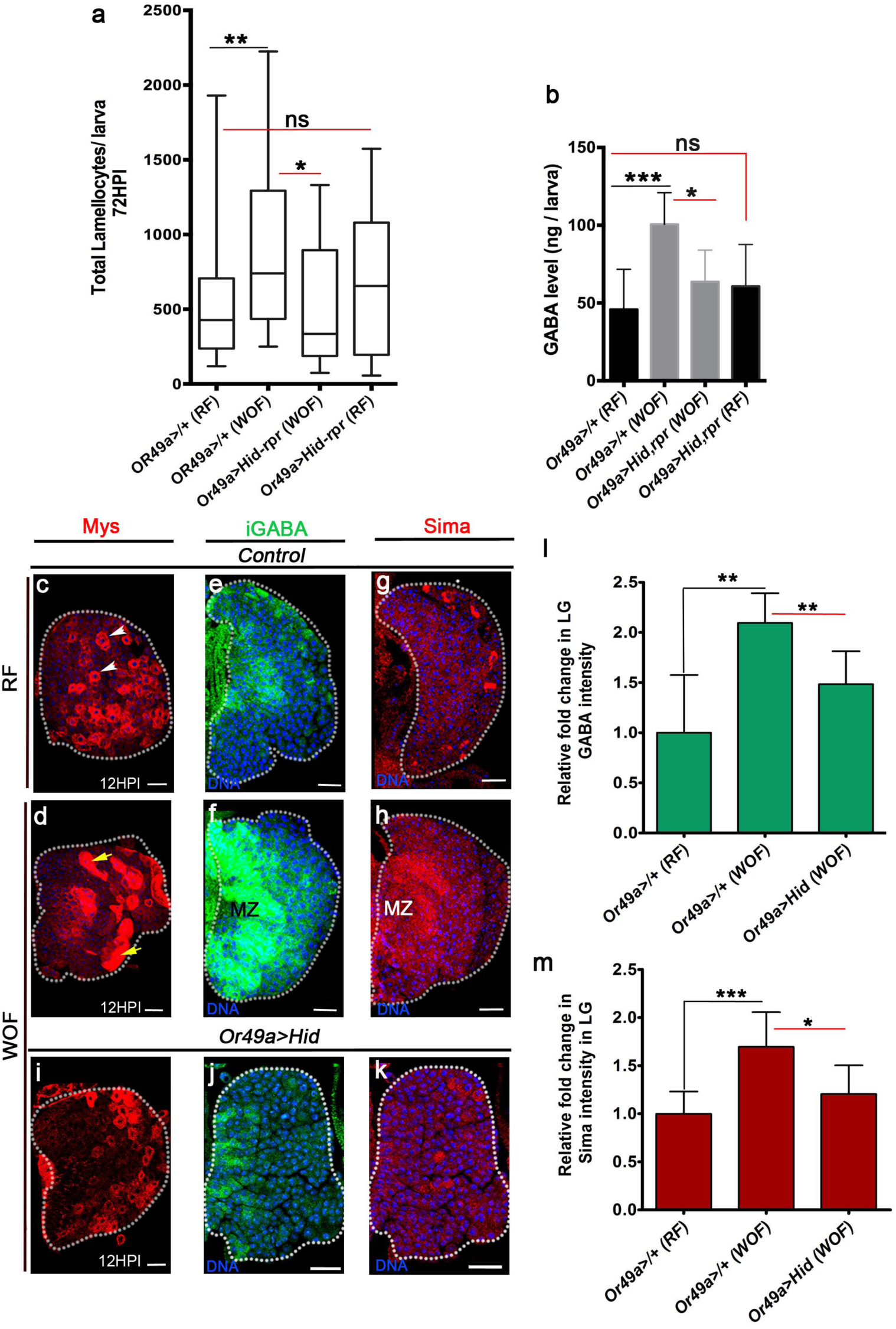
Physiological role for odors in blood cell immunity. DNA stained with DAPI (blue), iGABA (green), Sima protein (red). Scale bars = 20μm. RF is regular food, WOF is <u>w</u>asp <u>o</u>dor <u>f</u>ood and HPI indicates <u>h</u>ours <u>p</u>ost wasp-infection and ns is non-significant. In **a** median is shown is box plots and vertical bars represent the upper and lowest cell-counts. In panels **b** Mean with standard deviation is shown and in panels **l, m** fold change with standard error of mean (SEM) is shown. Statistical analysis, in panel **a** Mann-Whitney test, two-tailed and in panels **b, l, m** is un-paired *t*-test, two-tailed. “n” represents total number of larvae analyzed. **(a)** Quantification of total circulating lamellocyte numbers per larvae in controls (*Or49a>/+*) raised on RF (n=29), WOF (n=27, **p=0.009), *Or49a-Gal4, UAS-Hid, rpr* on WOF (n=18, *p= 0.03 compared to corresponding control on WOF) and *Or49a-Gal4, UAS-Hid, rpr* in RF (n=17). **(b)** Quantifications of hemolymph GABA from 3^rd^ instar larvae on RF (*Or49a>/+*), WOF (*Or49a>/+*, ***p=0.0008), *Or49a-Gal4, UAS-Hid, rpr* on WOF (*p=0.01) and *Or49a-Gal4, UAS-Hid, rpr* on RF. **(c** and **d)** Lamellocytes (red) detected at 12HPI in lymph glands from (**c**) RF (*Or49a>/+*) and (**d)** WOF (*Or49a>/+*) animals. WOF condition shows enlarged lamellocytes (red, yellow arrows) compared to smaller cells detected in RF (white arrowheads). **(e-h)** WOF condition leads to elevated blood cell iGABA level and Sima protein expression. Compare representative lymph gland images depicting (**e, f**) iGABA levels in (**e**) in RF (*Or49a>/+*) and (**f**) WOF (*Or49a>/+*) and (**g, h**) Sima protein in (**g**) RF and (**h**) WOF. WOF condition shows elevated expression of both iGABA and Sima, especially within the progenitor-cell compartment (Medullary Zone, MZ). See quantifications of relative fold change in iGABA and Sima intensities in **l, m** and Supplemental Fig. 10a for mean intensity values. (**i-k**) Lymph glands obtained from *Or49a-Gal4, UAS-Hid* animals reared in WOF fail to (**i**) show the lamellocyte response seen in WOF (compared with controls in **d**), (**j**) raise iGABA (compared with **f**) or (**k**) Sima protein (compare with **h**). The levels detected in *Or49a-Gal4, UAS-Hid* animals reared in WOF are comparable to levels seen in corresponding controls on RF (**c, e** and **g**). (**l, m**) Relative fold change in lymph gland intensities of (**l**) iGABA RF (*Or49a>/+*, n=8), WOF (*Or49a>/+*, n=6, **p=0.001), *Or49a>Hid* on WOF (n=8, **p=0.004) and (**m**) Sima protein on RF (n=8), WOF (n=8, ***p=0.0004), *Or49a>Hid* on WOF (n=8, *p=0.01). Corresponding mean intensities are shown in Supplemental Fig. 10a.

If wasp-preconditioning leads to improved lamellocytes formation at the expense of other immune cells remains unclear, but lymph gland development and blood cell profile of WOF animals were comparable to controls in regular food condition (Supplemental Fig. 8, h-l) although a minor (10%) increase in progenitor population was apparent (Supplemental Fig. 8, i and j). Circulating blood cell numbers remained unchanged (Supplemental Fig. 8l) and ectopic lamellocyte formation was barely detectable in these animals (See un-injured response in Supplemental Fig. 8, m-n). Thus, the heightened lamellocyte response in WOF animals most likely weren’t a consequence of any prior increase in blood cell numbers or lamellocyte differentiation. Apart from wasp infection, general injury responses in WOF animals also demonstrated a specific increase in lamellocyte potential only (Supplemental Fig. 8, m,n), without altering differentiation into other blood lineages (Supplemental Fig. 8, o-q).

Thus far, the data are consistent with a role for general olfactory input in maintaining immune competency. When olfactory activity is reduced (through ORNs killing, Orco mutation, Or42a mutation), immune competency is compromised. The experiments using wasp odor food also show that this odor-sensing axis further elevates the immune-competitiveness of the animals. If prior sensing of wasps provided the larvae with an immune adaptive value that is specific to the wasp odor and not a general consequence of exposure to elevated levels of odors leading to increased olfactory input over and above a basal threshold, was investigated. For this, we exposed *Drosophila* larvae from early development to odors of either attractive (acetic acid) or aversive (1-octen-3-ol and acetophenone) in nature. These experiments were conducted as undertaken for WOF (see methods for details). The pre-conditioned animals were followed by analysis of their lamellocyte differentiation potential in response to parasitic wasp-infection (Supplemental Fig. 9e). The odors were chosen based on their physiological relevance and their ability to induce strong behavioral responses (Kreher et al., 2005). Acetic acid activates Or42a (Kreher et al., 2005), 1-octen-3-ol is a fungal aversive odor and reported to influence *Drosophila* larval blood cell response (Inamdar and Bennett, 2014) and acetophenone is a well-established strong aversive odor (Kreher et al., 2005). We find diverse lamellocyte responses to varying odorants. Exposure to acetic acid or 1-octen-3-ol did not effect lamellocyte differentiation, however acetophenone led to a significant reduction in lamellocyte numbers (Supplemental Fig. 9e). Importantly, none of these odors manifested the lamellocyte increase seen with wasp-odors. Also, when assessed for blood cell iGABA levels, both acetic acid and 1-octen-3-ol did not reveal much difference (Supplemental Fig. 9 f-j). Acetophenone exposure however led to a stark reduction in iGABA levels which is consistent with the reduction in lamellocyte numbers detected in this condition. While these data reveal the diverse influence of varying odors on immune cells, they clearly show that increasing olfactory input over and above the basal threshold is not responsible for mediating the immune-phenotype seen in WOF. Also, not all aversive odors mediate the same immune response, thus highlighting specificity of odors in controlling immune phenotypes and that the sensing of predators/wasp odors in the environment indeed functions as an immune-adaptive component to prime superior lamellocyte potential. Next, the mechanism by which wasp-odors are sensed and signal to immune cells was investigated.

The detection of wasp odors in larvae is facilitated by activation of Or49a(Ebrahim et al., 2015). We found that ablation of Or49a (*Or49a>Hid*), in regular conditions did not manifest any reduction in lamellocyte numbers (Fig. 7a) or melanization response (Supplemental Fig. 10b). Neither did its loss reveal any difference in GABA (Supplemental Fig. 10d compared to c and i) or Sima protein (Supplemental Fig. 10g compared to j, m) expression. Contrastingly, in WOF condition, ablating Or49a significantly reduced the wasp-odor mediated increased lamellocyte numbers (Fig. 7a). Also, these animals did not demonstrate the increase in lamellocyte sizes detected in WOF controls (Fig. 7i compared to d). The increased hemolymph GABA levels seen in WOF conditions was no longer detected in *Or49a>Hid* WOF larvae either (Fig. 7b). Lymph gland iGABA and Sima levels in *Or49a>Hid* WOF animals was comparable to regular food conditions (Fig. 7, j, k compared to e, g) and did not show the increase seen in controls raised in WOF (Fig. 7, f, h and Supplemental Fig. 10a). Therefore, Or49a is not necessary for lamellocyte differentiation or regulation of basal GABA production and Sima expression. Unlike Or42a, where its loss led to significant reduction in basal GABA and Sima expression (Supplemental Fig 10 e, h and j), loss of Or49a did not show any difference. Thus implying, a requirement for Or49a only in wasp-odor condition to manifest the increase in GABA, Sima and immune potential, even though, it is not essential for the developmental control of GABA and Sima levels and the establishment of lamellocyte potential in blood cells. Genetic means to force activate either Or42a or Or49a under regular conditions also demonstrated that Or49a activation (*Or49a>TrpA1*) is sufficient to mediate an increase in lamellocyte numbers as seen in WOF condition (Supplemental Fig. 10k). This is undectable upon forced activation of Or42a (*Or42a>TrpA1*) (Supplemental Fig. 10l) and consistent with lamellocyte response seen upon enhanced acetic acid odor exposure (Supplemental Fig.9e).

Downstream of ORN stimulation, projection neuron (PN) activation is necessary for mediating the lamellocyte immune response (Supplemental Fig 7e). We therefore analyzed PN-Ca^2+^ reporter activity by employing a transcriptional reporter of induced intracellular Ca^2+^ (TRIC) that monitors slow neuronal responses over a developmental time course (Gao et al., 2015). In animals raised in wasp-odor condition, intense TRIC reporter activity is detected in a specific subset of projection neurons (Supplemental Fig. 10), which is undetectable in animals raised in either regular food or in acetic acid odor conditions. Implying, specific signals being transferred upon wasp odor detection to projection neurons that is not conducted by other odors. If this specific PN activation is allied to enhancing immune competency remains to be addressed.

In a physiological context we predict that as developing *Drosophila* larvae dwell into their food medium, sensing of food-related odors which are the predominant odors generally present in the environment leads to activation of the neuronal circuit (ORN-PN-Kurs6^+^GABA^+^ neuronal route) that stimulates Kurs6^+^GABA^+^ neurosecretory cells to release GABA. This establishes the non-autonomous metabolic axis to control blood progenitor Sima levels. In the incidence of wasp-infestation, the larvae rearing within this environment detect wasp-odors via Or49a and the combinatorial ORN signaling of Or42a and Or49a leads to activation of specific subset of PNs most likely responsible for enhancing Kurs6 GABA production and release (Fig. 8). The involvement of other ORs in immune response, and their function either independently or in combination cannot be ruled out.

**Figure 8.**
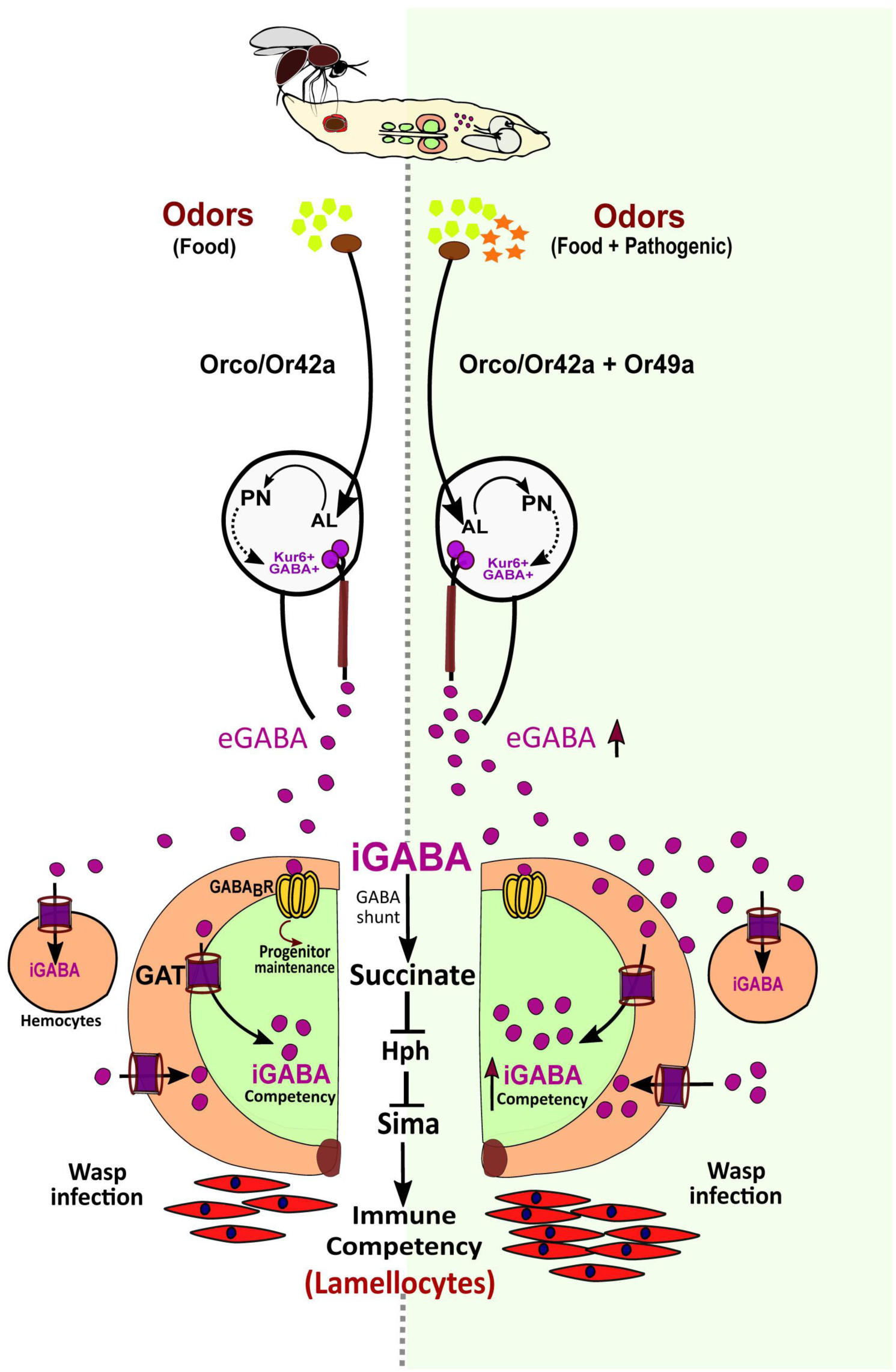
Developmental control of immune-competency by environmental odors. *Drosophila* larvae spend most of their time dwelling in food. The odors derived from this eco-system defines an integral immune-component during hematopoiesis. Sensing food via Or42a stimulates projection neurons (PN) leading to down-stream activation of Kurs6^+^GABA^+^ neurosecretory cells, which mediate release of GABA (eGABA) into the hemolymph. eGABA is internalized by blood cells via GABA-transporter (Gat) and its subsequent intracellular metabolism through the GABA-shunt pathway leads to stabilization of Sima protein in them and this establishes their immune-competency to differentiate into lamellocytes. Physiologically, this sensory odor axis is co-opted to detect environmental pathogenic wasp-odors. Upon detection of wasps via Or49a in the preconditioned media (WOF), the combinatorial stimulation of both Or42a and Or49a, elevates neuronal GABA release, leading to increase blood cell iGABA mediated Sima expression. This developmentally establishes superior immune-competency to withstand the immune-challenge.

## DISCUSSION

Systemic control of immune-cell fate specification and priming is not well-understood. In this study, we describe a fundamental role for olfaction in priming immune-competency that is necessary to respond to pathogenic insults (Fig. 8). We show that during *Drosophila* larval development, the detection of food-odors from their environment mediates the activation of neuronally-derived cues necessary to stimulate the systemic release of GABA from a subset of neurosecretory cells. Developmentally, GABA is sensed by blood cells both as a signaling ligand via GABA_B_R to promote blood-progenitor maintenance, and as a metabolite, where its internalization by blood cells via a functional transporter (Gat) establishes their immune-competency. iGABA catabolism through the GABA-shunt pathway prevents Sima protein from proteosomal mediated degradation. Sima is both necessary and sufficient for lamellocyte induction. Rearing *Drosophila* larvae in parasitoid-threatened environmental conditions, raises their systemic GABA levels, this consequently elevates the expression of blood-progenitor Sima levels. We find that supplementing wild-type animals with additional GABA or succinate also elevates blood progenitor Sima expression and lamellocyte responses (Supplemental Fig. 11, a-d) which is lost upon progenitor-cell specific (Dome^+^) abrogation of *sima* expression (Supplemental Fig. 11d). Limited GABA availability during development therefore restricts the extent of blood-progenitor Sima levels and consequently the lamellocyte induction capacity.

### Olfaction, GABA and blood development

The plasticity of the hematopoietic system in *Drosophila* larvae with olfactory dysfunction or abrogated GABA metabolism to support a cellular immune differentiation program that responds on demand is lost, even though blood cell differentiation in homeostasis is intact in these mutants. Demonstrating that immune-potential is independently controlled and not always a consequence of loss of progenitor maintenance. Thus, olfactory control of progenitor maintenance via GABA/GABA_B_R signalling alongside the use of GABA metabolism in immunity exemplifies the role of olfaction in blood development and immune-competency. In mammals loss of olfaction as in olfactory bulbectomized mice or humans with olfactory dysfunction, is associated with heightened inflammation and a failure to respond when challenged(Strous and Shoenfeld, 2006). While these observations are similar to what is described in this study and highlight a well-conserved phenomenon to use olfaction to control immunity, the broader implications of our work in higher model systems with more complex blood repertoires remains to be validated.

### Physiological role for the olfaction-immune axis

The physiological implications of the olfaction-immune axis is not well-understood. Priming cellular immune potential during development by employing environmental odors to stimulate a long-range metabolic cross-talk described here represents the first example highlighting its relevance in animal physiology. The use of this connection during *Drosophila* larval development to support an innate immune training component is indeed intriguing. Most often, animals in the wild are dwelling in surroundings with pathogenic threats in their environment. Such is also the case with *Drosophila* larvae in the wild where almost 80% of them are infected with wasps of this species(Fleury et al., 2004). Larvae being more vulnerable to infection with their limited abilities to avoid, an adaptive mechanism that enhances immunity poses a viable option to withstand such challenges. The control of inflammatory response by olfactory cues may therefore have arisen as a means to deal with unfavourable conditions. The use of general broad-odors to establish basic immune-potency that can be further manipulated depending on environmental conditions, exemplifies a rheostat-like control by the olfactory axis. Such impact of odor-experience as a direct handle into fine-tuning immune metabolism to enable an immune advantage is the first *in vivo* description of its kind. In our analysis we have tested one attractive and three diferent aversive odors. First, we find that over and above exposure to odors did not mediate the same affect on larval immune responses to wasp-infection. Indicating it is not the overall strength of the olfactory input. Second, excessive exposure to attractive odors did not affect any aspect of blood development or immune response while minimal food odor condition severely down-regulates lamellocyte differentiation. Thus a certain threshold of attractive odors is necessary but beyond this the system does not respond. Exposure to a fungal aversive odorant, 1-octen-3-ol, shown to affect larval plasmatocyte responses by controlling the levels of nitric oxide signaling(Inamdar and Bennett, 2014), showed no dramatic difference to wasp-infection driven lamellocyte differentiation or GABA levels. These data highlight diverse systemic modules employed by odors to manipulate immune cells with specificity. Contrary to this, detection of pathogenic wasp odors (via Or49a) co-opts the olfaction-GABA axis to further enhance neuronal GABA levels and subsequently their immune-competency to generate more lamellocytes. This phenomenon enables an innate adaptive component which the animal may be benefitted from when thriving in wasp prone environments often encountered in the wild (Fig. 8). Exposure to acetophenone, a yet another aversive odor although unclear of its physiology, demonstrates a rather opposite phenotype that leads to down-regulation of basal GABA production and immune response.

### ORN-PN crosstalk and immunity

OR42a being the most pre-dominant OR is activated in response to food-related odors. This establishes the route that systemically connects the olfactory modality to the development of the immune system (ORN-PN-Kurs6-GABA-hematopoietic cells). Detection of pathogenic odors via Or49a also co-opts the olfaction-GABA axis to further raise GABA levels and subsequently their immune-competency (Fig. 8). The connection established by sensing of food odors therefore sensitizes the animal to respond to wasps with heightened efficiency. Genetically activating Or49a can also reciprocate a similar enhancement of immune response as seen in wasp-odor enriched conditions. However, activation of Or42a (*Or42a>TrpA1*) failed to increase lamellocyte responses any further. Implying varying potentials of ORNs and their underlying downstream regulation. Although not conclusive, an elevated projection neuron Ca^2+^ reporter activity in animals raised in wasp-odor conditions compared to regular conditions, lends us to hypothesize specific activation of projection neurons as a key regulator of the olfactory circuit leading to heightened stimulation of neuronal GABA production upon wasp-odor exposure. Cross-talk between individual ORN and their respective glomeruli and how they control PN activity (Berck et al., 2016) remains to be explored and awaits further investigation.

Being a key pro-survival sensory modality, this study expands our current understanding of olfaction beyond modulation of animal behaviour, implying more diverse physiological contexts(Riera et al., 2017) than previously known. Both GABA_B_R and Gat are detected in immune cells of myeloid(Stuckey et al., 2005) and lymphoid origin(Jin et al., 2013). Common myeloid progenitor cells maintain elevated ROS(Tothova et al., 2007), in a HIFα dependent manner (Cramer et al., 2003) and for priming immunity(Cheng et al., 2014). Moreover, stimulated macrophages shift to GABA as their metabolic resource to generate succinate and mediate inflammatory response via HIFα (Tannahill et al., 2013). These are some commonalities between the mammalian myeloid system and *Drosophila* hematopoietic system that strengthen the importance of the work presented here. It will therefore be interesting to determine, if elements of the olfaction/immune axis described here are relevant for general myeloid development and competency. Not only an understanding of how immunity is controlled by odors will emerge, these studies will also yield better insights into employing olfactory routes to train immunity in development and disease.

## METHODS

### *Drosophila* husbandry, stocks, genetics and wasp infections

The following *Drosophila* stocks were used in this study: *w^1118^* (wild type*, wt*) *DomeMESO-Gal4, UAS-EYFP* (Banerjee lab)*, Hml^Δ^-Gal4, UAS-2xEGFP* (S.Sinenko)*, Kurs6-Gal4* (G. Korge)*, Orco-gal4* (BL 26818)*, Or49a-Gal4* (BL 9985)*, Or42a-Gal4* (BL 9969)*, GH146-Gal4*(Shim et al., 2013)), *orco^1^* (BL 23129 (Shim et al., 2013))*, UAS-TRIC* (BL 61681(Gao et al., 2015)), *UAS-Hid, rpr/CyoGFP, UAS-Hid, UAS-TrpA1* (BL 26263)*, UAS-sima* (P.Wappner)*, UAS-Ldh* (Flyorf F002924), *UAS-Hph* (C.Frei) and *gabat^PL00338^* (A.Sehgal), TRIC (BL 61680 (Gao et al., 2015)). The *RNAi* stocks were obtained either from Vienna (VDRC) or Bloomington (BL) *Drosophila* stock centers. The lines used for the study are: *Gad1^RNAi^* (BL 28079 (Shim et al., 2013))*, GABA_B_R1^RNAi^* (BL 28353 (Shim et al., 2013)) *GABA_B_R2^RNAi^* (BL27699 (Shim et al., 2013)), *Gat^RNAi^* (BL 29422 (Stork et al., 2014))*, Gabat^RNAi^* (VDRC 110468KK) *Ssadh^RNAi^* (VDRC 106637)*, CG3379^RNAi^ (α-KDH*, BL 34101)*, skap^RNAi^* (BL 55168), *SdhA^RNAi^* (VDRC 330053), *sima^RNAi^* (HMS00832 (Wang et al., 2016))*, Hph^RNAi^* (VDRC 103382 (Mukherjee et al., 2011)*), Notch^DN^* (Mukherjee et al., 2011)*, Notch^RNAi^* (VDRC, v1112 (Liu et al., 2017))*, Ldh^RNAi^* (BL33640 (Li et al., 2017))*, Tgo^RNAi^* (BL 26740, VDRC 103382 (Mukherjee et al., 2011)), *ChAT^RNAi^* (BL25856 (Shim et al., 2013)). All fly stocks were reared on corn meal agar food medium with yeast supplementation at 25°C incubator unless specified. The crosses involving RNA*i* lines were maintained at 29°C to maximize the efficacy of the *Gal4/UAS* RNA*i* system. Controls correspond to either *w^1118^* (wild type) or Gal4 drivers crossed with *w^1118^*. All the RNA*i* stocks were tested for their knockdown efficiencies by using a ubiquitous driver to express these lines followed by isolation of total mRNA from whole animals subjecting them to qRT-PCR analysis with respective primers. *RNAi* knockdown efficiencies of the respective lines are: *Gat^RNAi^* (97.7%), *CG3379^RNAi^ (α-KDH*, 95%), *Ssadh^RNAi^* (45%) and *skap^RNAi^* (40%).

To control for backgrounds and off target affects, the RNA*i* lines and genetic rescue combinations were tested by crossing them to *w^1118^* and assessed for lamellocyte formation. All the lines tested made lamellocytes comparable to *w^1118^* (Supplemental Fig. 11e).

*Leptopilina boulardi* were maintained and the infections were set up as previously described (Schlenke et al., 2007). Briefly, fly larvae aged 60 ± 2 hours after egg laying were exposed to 8-10 female and 3-4 male wasps for 6-8 h. Infected larvae with a visible poke were collected for further experimental analysis.

### Immunostaining and immunohistochemistry

For staining circulating cells, 3^rd^ instar larvae were collected and washed in 1X PBS and transferred to Teflon coated slides (Immuno-Cell #2015 C 30) followed by staining protocol previously described (Jung *et al.*, 2005). Lymph glands isolated from larvae were also stained following the similar staining protocol. Immunohistochemistry on lymph gland and circulating blood cells was performed with the following primary antibodies: mouse αP1 (1:100, I. Ando), rabbit αPxn (1:2000, J. Shim), mouse αLz (1:100, DSHB), rabbit αPPO (1:1000) mouse αHnt (1:100, DSHB), rabbit αGABA (1:100, Sigma), mouse αWg (1:5, DSHB 4D4), rat αCi (1:5, DSHB 2A1), rabbit αphospho-CaMKII (1:100, Cell Signaling #3361), αGat (M. Freeman), guinea pig αSima (1:100, U. Banerjee). The following secondary antibodies were used at 1:500 dilutions: FITC, Cy3 and Cy5 (Jackson Immuno Research Laboratories). Phalloidin (Sigma-Aldrich # 94072) was used at 1:100 dilution to stain cell morphologies and nuclei were visualized using DAPI. Samples were mounted with Vectashield (Vector Laboratories). A minimum of five independent biological replicates were analyzed from which one representative image is shown.

### *In situ* hybridization

Digoxigenin (DIG)-labelled probes for *in situ* hybridization was synthesized by PCR using DIG RNA labelling kit (Roche #11175025910). The probes for *Ssadh* and *CG33791* (*αKDH*) genes were generated using primers mentioned previously that were fused to a T7 promoter sequence. Finally the probes were applied to dissected lymph gland tissues prepared for hybridization following the previously published protocol (Shim et al., 2012).

### Quantitative Real-Time PCR analysis

Total RNA was extracted using Trizol reagent (Invitrogen, USA). For lymph gland analysis, RNA was obtained from 3^rd^ instars (#150 for each genotype). The total RNA extracted was reverse transcribed with Super Mix kit (Invitrogen) and followed by quantitative real-time PCR (QPCR) with SYBR Green PCR master mix kit (Applied Biosystems). The relative expression was normalized against *rp49* gene. The respective primers used are the following:

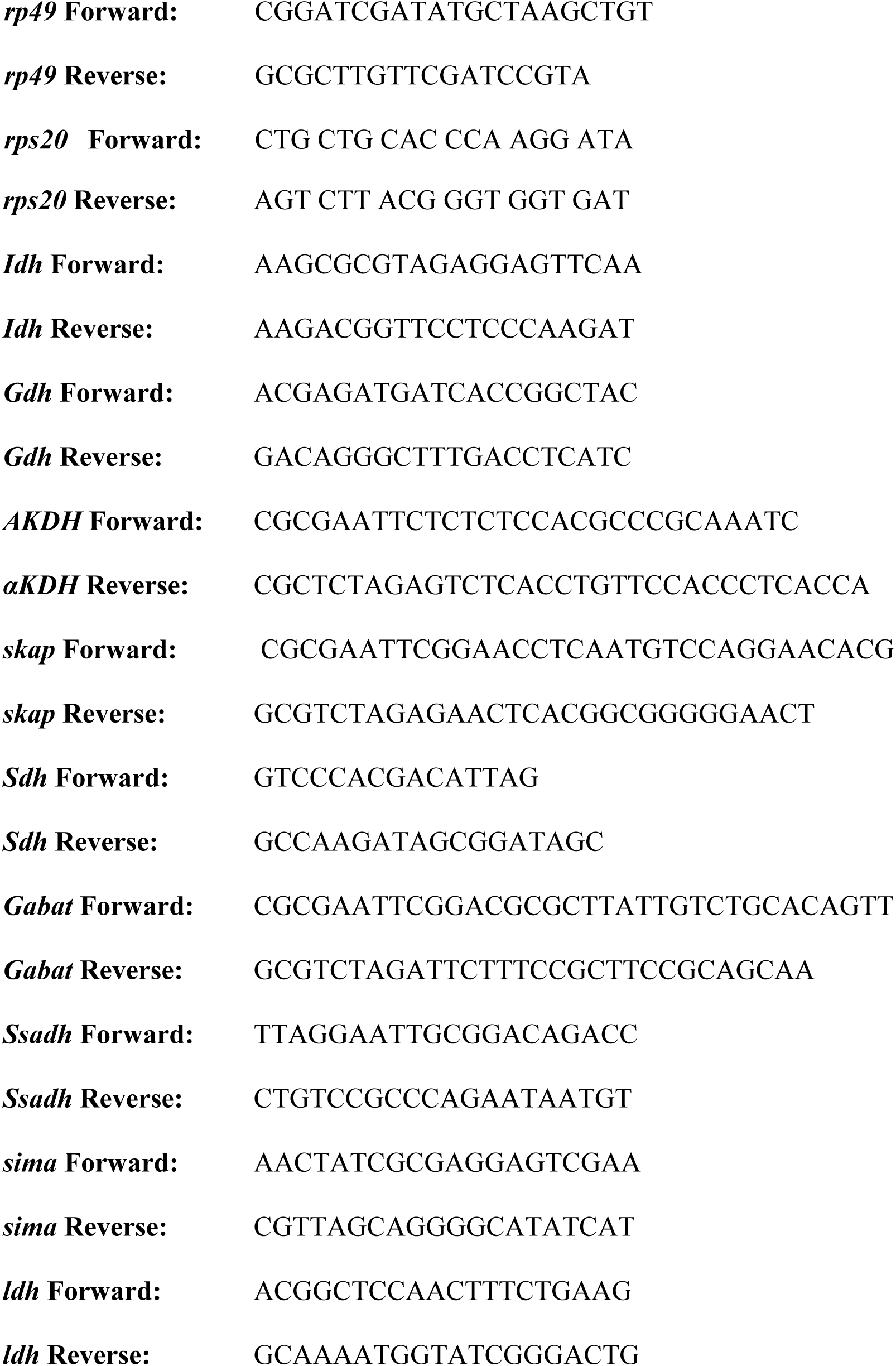

### Imaging

Immuno-stained images were acquired using Olympus FV1000 confocal microscopy or Nikon C2 Si-plus system under a 20X air or 40X oil-immersion objective. Bright field images were obtained on OlympusSZ10 or Zeiss Axiocam.

### Hemocyte quantification

Lamellocyte population was quantified both in the lymph gland and in circulation. For circulating lamellocyte counts, individual larva were bled per well and the cells were manually counted. Lamellocytes were detected by their large flattened morphology visualized by using Phalloidin counterstain as previously published or Myospheroid/L4(Anderl et al., 2016; Small et al., 2014,). For counting lamellocytes in lymph glands, the tissues were counter-stained with phalloidin or Myospheroid and imaged to obtain Z-stacks. Lamellocytes across Z-stacks were counted per lobe. To calculate total blood cell counts/mm^2^, five images covering the field of view with hemocytes were acquired per well. The magnification was kept constant at 20X. The hemocytes in these views were then counted using the ImageJ software plugin Analyze particles tool. Circulating cell numbers obtained are quantified per larvae. In all experiments, control genotypes were analyzed in parallel to the experimental tests. For each experiment, a minimum of 3-5 biological replicates were analyzed. This was repeated independently at least two times and the quantifications represent the median or the mean of all the biological replicates.

### Quantification of lymph gland phenotypes

All images were quantified using ImageJ software. Lymph gland area analysis was done as described in (Shim et al., 2012). Roughly, middle three confocal Z-stacks were merged and thresholded, selected and area was measured. This was done for respective zones and the area is represented in percent values. Controls were analysed in parallel to the tests every time. A minimum of 5 animals were analysed each time and the experiment was repeated at least three times. The quantifications represent the mean of the three independent experimental sets.

For quantifying mean intensities in lymph gland tissues it was calculated as described in literature (Louradour et al., 2017),(Morin-Poulard et al., 2016). Briefly, the relevant stacks of the lymph gland images were selected, the area to be measured per lobe was defined using the select tool and was then thresholded followed by intensity measures. The relative fold change in intensities per lobe was calculated using mean intensity values. For all intensity quantifications, the laser setting for each individual experimental set-up was kept constant. Controls were analysed in parallel to the tests every time. A minimum of 5 animals were analysed each time and the experiment was repeated at least three times. The quantifications shown are for one set.

### Melanization

To score “% melanized larvae”, wasp-infected *Drosophila* larvae obtained at 96 hours post-infection were first scored under the dissecting microscope for black spots in the cuticle and then dissected to identify *D. melanogaster* larvae with melanized wasp-eggs as described earlier (Yang et al., 2015). For each trial, the percentage of infected larvae with visible melanization was quantified. Compared to larvae without any melanized wasp-eggs, larvae with melanization response were scored as positive and the respective percentage of positive larvae in each cohort was calculated. A 100% melanization percentage would indicate that all of the infected *D. melanogaster* larvae showed visible melanization; while a value of 0% would indicate that none of the infected *D. melanogaster* larvae showed visible melanization.

To score “melanization efficiencies per larvae”, individual *Drosophila* larvae at 72 hours post infection were dissected to obtain the deposited wasp-eggs. The number of un-melanized and melanized wasp-eggs per larvae were counted and the percent melanization efficiency/larvae calculated by dividing the number of melanized wasp-eggs to the total number of wasp-eggs.

In each independent experimental set, at least 10 animals per genotype were analyzed and this was repeated a minimum of 3 times. The graphical quantifications represent the percentages of total melanized and un-melanized larvae obtained from all the independent experimental sets.

### GABA measurements

GABA measures in circulation were conducted by bleeding five wandering third-instar larvae to extract their hemolymph as previously published (Shim et al., 2013) and analyzed using LC-MS/SRM method (Agilent 1290 Infinity UHPLC). This was done for minimum of 15 larvae per genotype and repeated three times. The quantifications shown represents the mean of all the repeats.

### Minimal odor and odor infused food preparation

The minimal odor food was prepared as described previously (Shim et al., 2013). For wasp odor food, it was prepared by placing sealed dialysis tubing with low molecular weight cut-off (Spectra/Por Dialysis tubing MWCO 500-1000 D) containing *L. boulardi* wasps in the proportion of 15 females and 8 males into regular food medium. This set up allows odorant cues to pass through without diffusion of any macromolecular substance. The food was freshly prepared each time.

For exposure to others odorant (acetic acid, 1-octen-3-ol and acetophenone) larvae were treated with respective odorants of the highest available purity (>99% from Sigma) Acetic acid (Sigma 2722), 1-octen-3-ol (Sigma O5284 ) acetophenone (Sigma 00790) at concentrations as previously described in (Kreher et al., 2005). These odorants were constituted in Mineral oil (Sigma M5904) to obtain 10^−2^ dilution. 40 µl of diluted odorant was placed on whatman filter paper, which was then placed inside the vial containing regular fly-food media. Every 18-24 hr 40ul of odorants were again infused onto the same whatman paper to provide constant exposure of odors into the food.

*Drosophila* larvae from 1^st^ instar stage were reared in the different odorant infused food media until wandering 3^rd^ instar stage. To quantify the influence of odors in immune response, multiple *controls* were tested on regular and odor infused food. In each experimental set a minimum of 10 larvae analyzed. This was repeated a minimum of three times for every genotype.

### Succinate and GABA supplementation

Succinate (Sodium succinate dibasic hexahydrate, Sigma #SLBM6312V) and GABA (Sigma #BCBR7698V) enriched diets were prepared by supplementing regular fly food with respective amounts by weight/volume measures of succinate or GABA to achieve 3% or 5% concentrations respectively. Eggs were collected in these supplemented diets and reared until analysis of their respective tissues.

### Statistical analyses

All statistical analyses was performed using GraphPad Prism five software and Microsoft Excel 2010. The medians were analyzed using Mann-Whitney test, two-tailed and means were analyzed with unpaired *t*-test, two-tailed.

## ACKNOWLEDGEMENTS

We thank U. Banerjee for Sima antibody, M. Freeman for Gat antibody, A. Sehgal for *gabat^PL00338^*, Shannon Olsson for odor experiments, N. Mortemier and T. Schlenke for *L.boulardi* stock and VDRC (Austria), Flybase and the Bloomington Stock Center for fly stocks, NCBS, CCAMP for their fly, imaging and metabolomics facilities. We thank Apurva Sarin and inStem collegues for helpful discussion and comments on the manuscript. Due to space limitations, we apologize to our colleagues whose work is not cited. This study was supported by the DBT-Center of Excellence grant BT/PR13446/COE/34/30/2015, DST-ECR ECR/2015/000390 and DBT Ramalingaswami Re-entry Fellowship to T.M and Basic Science Research Program through National Research Foundation (NRF-2014S1A2A2028388 and NRF-2017R1C1B2007343) to J.S. S.M is a Graduate student at inStem, in the Mukherjee lab.

## Supplemental Data

**Supplemental Figure 1.**
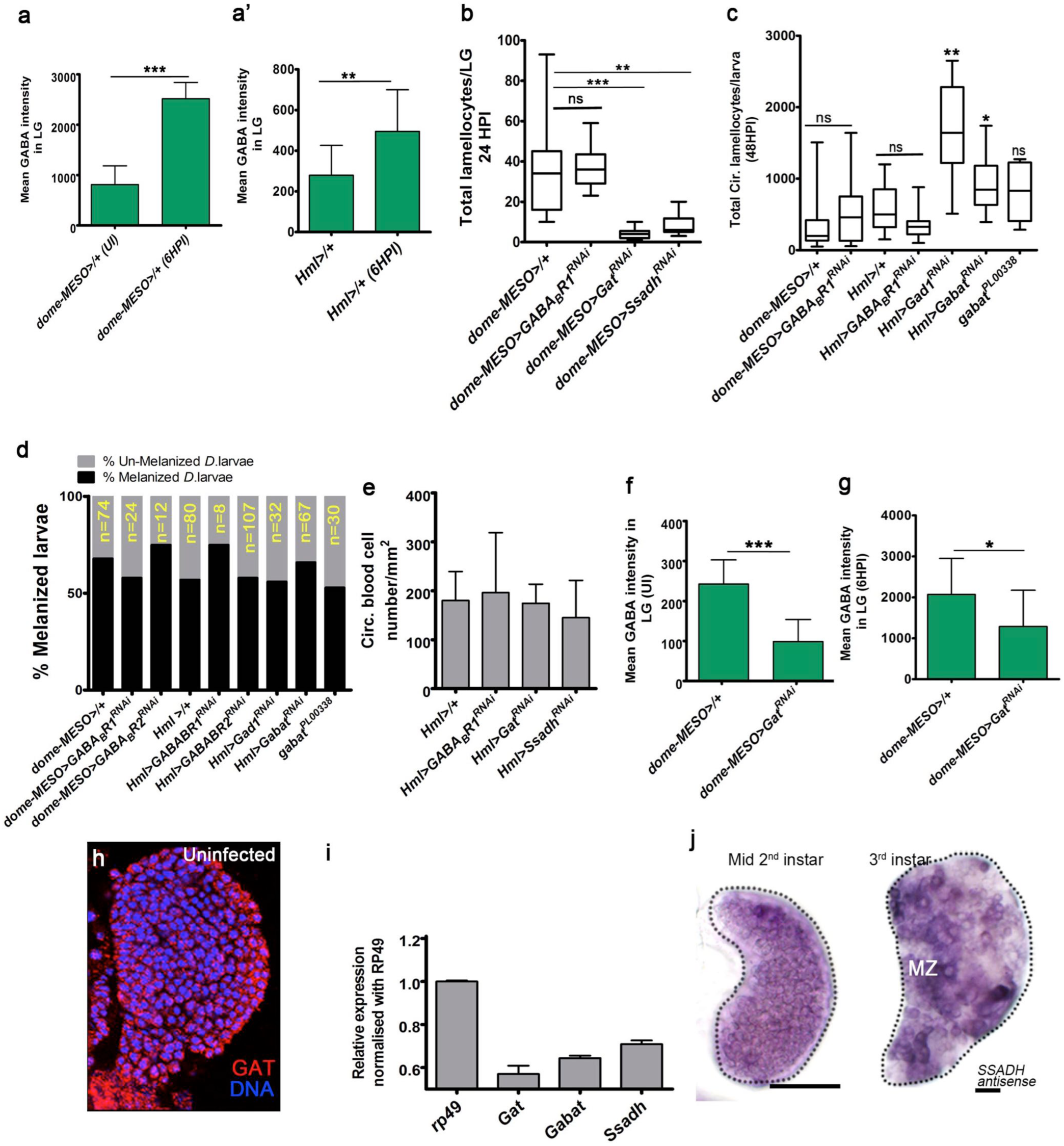
Distinctive roles for the GABA-shunt pathway in lamellocyte differentiation. UI is un-infected, HPI is <u>h</u>ours <u>p</u>ost wasp-infection, LG is lymph gland, ns is non-significant. Graphs in panel **a, a’, e, f, g, i** represents Mean ± Standard deviation (Mean ± SD), in **b, c** median is shown in box plots and vertical bars represent upper and lowest cell counts. Statistical analysis in panels **a, a’, e, f** and **g** is un-paired *t*-test, two-tailed and in panels **b, c** is Mann-Whitney, two-tailed. “n” represents total number of larvae analyzed. (**a, a’**) Mean intensity plot of iGABA expression in controls lymph glands from (**a**) *domeMESO-Gal4, UAS-GFP/+* in un-infected (n= 10, 812 ± 373) and at 6HPI (n=11, 2516 ± 324, ***p<0.0001) and (**a’**) *Hml^Δ^-Gal4, UAS-GFP/+* in un-infected (n= 11, 279.4 ± 146) and at 6HPI (n=12, 495 ± 205, **p= 0.0091). **(b)** Total lamellocyte quantification per lymph gland. *domeMESO-Gal4, UAS-GFP/+* (control, n=15), *domeMESO-Gal4, UAS-GFP; UAS-GABA_B_R1^RNAi^* (n= 17), *domeMESO-Gal4, UAS-GFP; UAS-Gat^RNAi^* (n= 13, ***p<0.0001) and *domeMESO-Gal4, UAS-GFP; UAS-Ssadh^RNAi^* (n= 8, **p=0.001). **(c)** Total circulating lamellocyte quantification per larvae. *domeMESO-Gal4, UAS-GFP/+* (control, n=30), *domeMESO-Gal4, UAS-GFP; UAS-GABA_B_R1^RNAi^* (n= 24), *Hml^Δ^-Gal4, UAS-GFP/+* (n= 9), *Hml^Δ^-Gal4, UAS-GFP; UAS-GABA_B_R1^RNAi^* (n=12), *Hml^Δ^-Gal4, UAS-GFP; UAS-Gad1^RNAi^* (n= 7), *Hml^Δ^-Gal4, UAS-GFP; UAS-Gabat^RNAi^* (n= 13) and *gabat^PL00338^* (n=9). **(d)** Graphical representation of melanization response. Also see Table S1. **(e)** Quantification of blood-cell densities represented as circulating blood cell count/mm^2^ of the respective genotypes. Refer to Table S3 for absolute values. **(f)** Mean intensity plot of iGABA expression in lymph glands from un-infected *domeMESO-Gal4, UAS-GFP/+* (control, n=10, 243 ± 60.2) and *domeMESO-Gal4, UAS-GFP; UAS-Gat^RNAi^* (n= 10, 99 ± 55.1, ***p<0.0001). **(g)** Mean intensity plot of iGABA expression in lymph glands at 6HPI in *domeMESO-Gal4, UAS-GFP/+* (control, n=10, 2069 ± 880.3) and *domeMESO-Gal4, UAS-GFP; UAS-Gat^RNAi^* (n= 13, 1287 ± 887, *p=0.047). **(h)** Lymph glands from uninfected larvae showing uniform Gat (red) expression in them. DNA marked with DAPI (blue). **(i)** Relative *mRNA* expression of *Gat* and GABA-shunt enzymes in RNA extracted from 3^rd^ instar wild-type larval lymph glands. **(j)** Representative *in situ* hybridization images showing *Ssadh mRNA* expression in lymph glands from mid-2^nd^ and 3^rd^ instar control larva. Uniform expression is detected in all blood cells in early time-points. Later the expression is comparatively elevated within the prospective progenitor compartment (medullary zone, MZ).

**Supplemental Figure 2.**
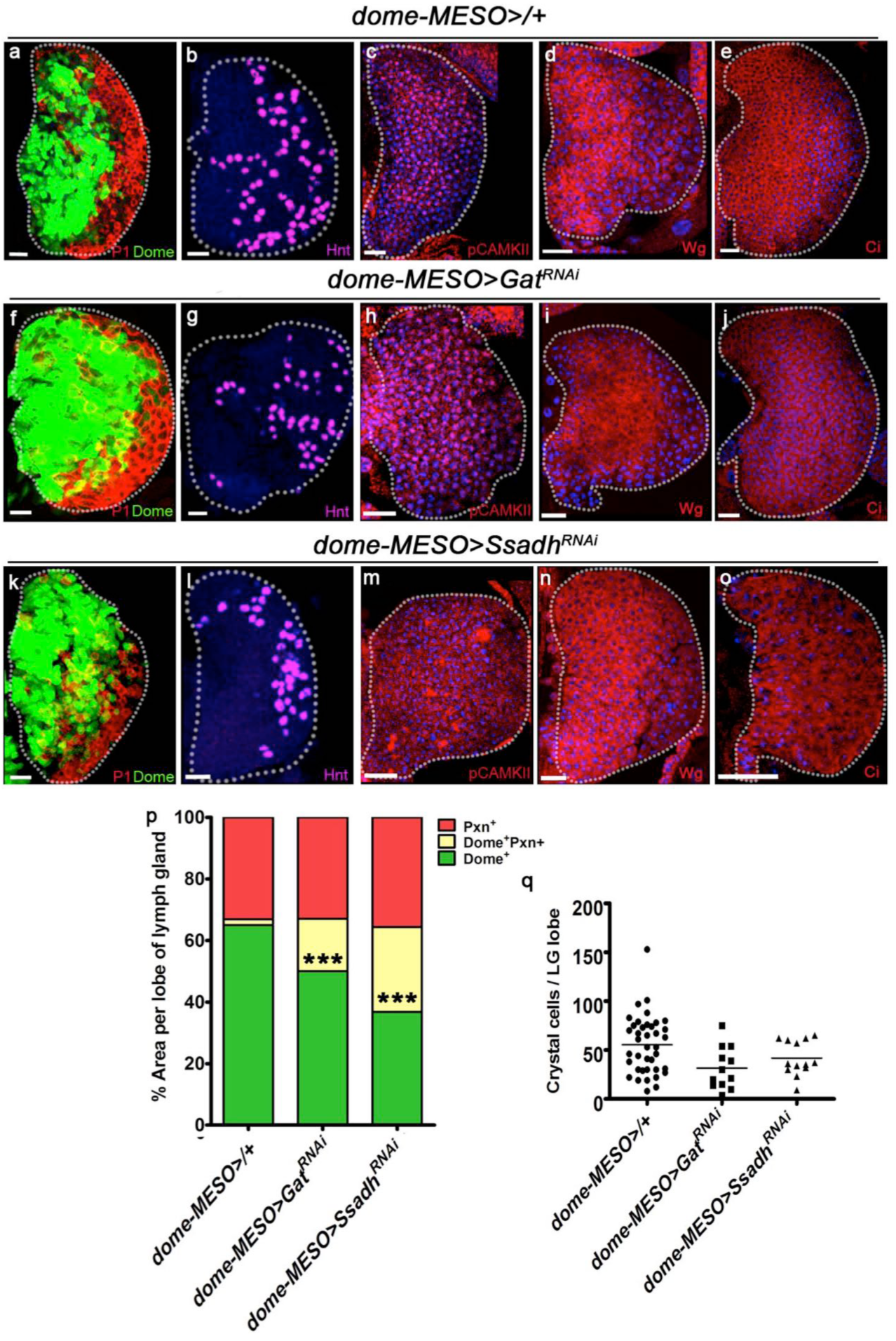
GABA-shunt pathway is dispensable for normal hematopoiesis. DNA is stained with DAPI (blue). Representative 3^rd^ instar lymph gland images showing progenitors in green (Domeless, Dome), plasmatocytes in red (P1) and crystal cells in magenta (Hindsight, Hnt). Scale bar = 20μm and “n” is total number of larvae analyzed. (**a-e**) Control (*domeMESO-Gal4, UAS-GFP/+*) lymph glands showing **(a)** Dome (green), P1 (red)), (**b**) Hnt (magenta), (**c**) pCAMPKII (red), (**d**) Wingless (Wg, red) and (**e**) Cubitous interruptus (Ci, red) expression. (**f-o**) Compared to control in **a-e**, (**f-j**) *Gat^RNAi^* (*domeMESO-Gal4, UAS-GFP; UAS-Gat^RNAi^*) or (**k-o**) *Ssadh^RNAi^* (*domeMESO-Gal4, UAS-GFP; UAS-Ssadh^RNAi^*) show comparable expression of **(f, k)** Dome (green), P1 (red), (**g, l**) Hnt (magenta), (**h, m**) pCAMPKII (red), (**i, n**) Wingless (Wg, red) and (**j, o**) Cubitous interruptus (Ci, red). (**p**) Graphical representation of percentage areas of progenitor cells (green, Dome^+^ only), intermediate population (IP, yellow, Dome^+^ Pxn^+^) and differentiating blood cells (red, Peroxidasin^+^ only). An expansion of IP is noted in genetic knock-downs of *Gat* and *Ssadh* (***p<0.0001 in comparison to control *domeMESO-Gal4, UAS-GFP/+*). See Table S2. (**q**) Quantification of total crystal cells per lymph gland lobe in control (*domeMESO-Gal4, UAS-GFP/+*, n= 20), *Gat^RNAi^* (*domeMESO-Gal4, UAS-GFP; UAS-Gat ^RNAi^*, n=6) and *Ssadh^RNAi^* (*domeMESO-Gal4, UAS-GFP; UAS-Ssadh ^RNAi^*, n=7). Also see Table S4.

**Supplemental Figure 3.**
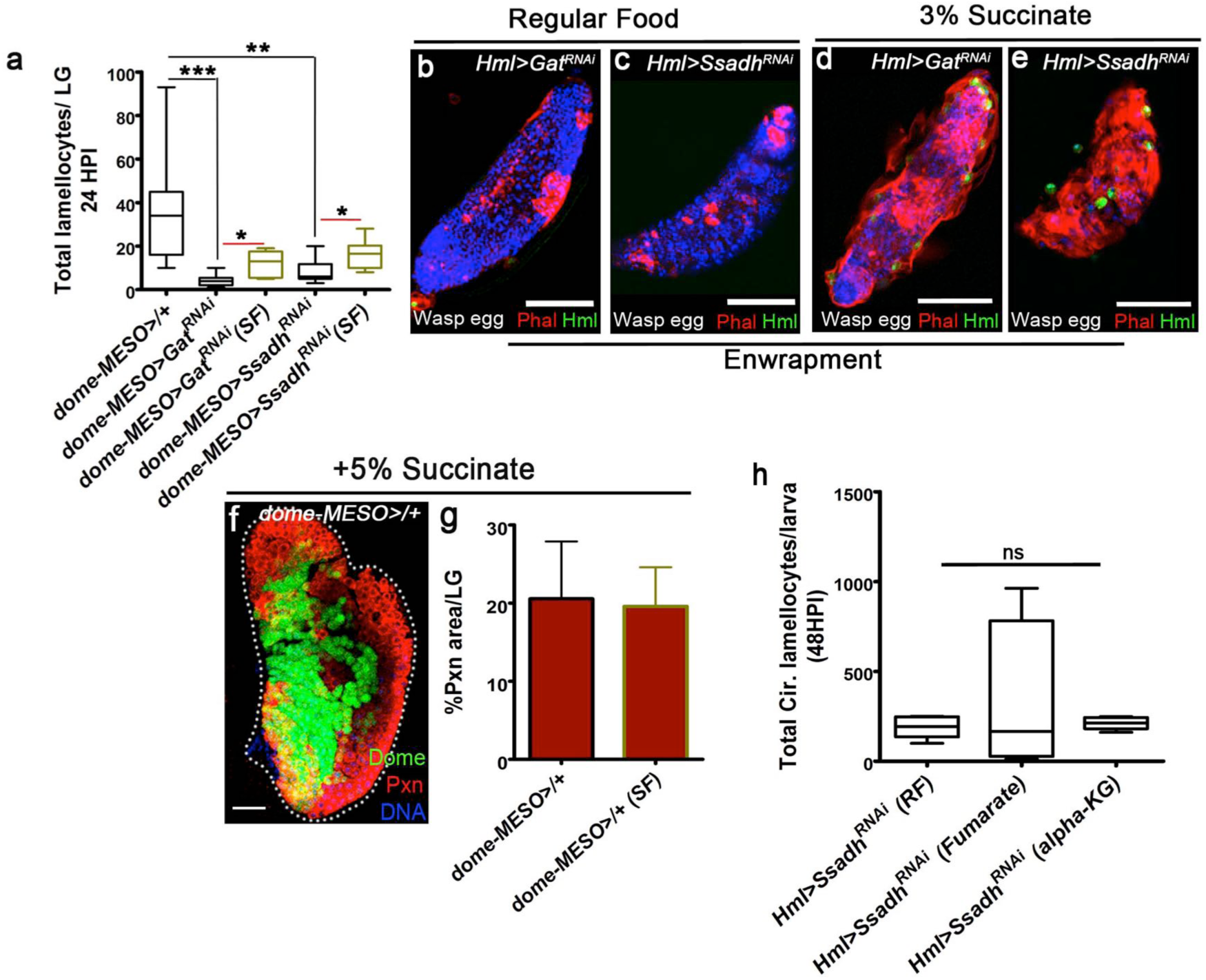
GABA-shunt derived succinate is necessary for lamellocyte differentiation. Nuclei are marked with DAPI (blue). Phalloidin (red) marks all blood cells and lamellocytes are characterized by their large flattened morphology. Scale bars = 100μm. HPI is <u>h</u>ours <u>p</u>ost wasp-infection, SF is succinate food, RF is regular food, ns is non-significant. In panels **a, h** median is shown in box plots and vertical bars represent upper and lowest cell counts. In **g**, Mean ± Standard deviation (Mean ± SD) is shown. Statistical analysis in panel **a** and **h** is Mann-Whitney test, two-tailed and in **g** is un-paired *t*-test, two-tailed. “n” is total number of larvae analyzed. (**a**) Quantifications of total lamellocytes in lymph glands of *domeMESO-Gal4, UAS-GFP/+* on RF (control, n=15), *domeMESO-Gal4, UAS-GFP; UAS-Gat^RNAi^* on RF (n=13, ***p<0.0001), *domeMESO-Gal4, UAS-GFP; UAS-Gat^RNAi^* on SF (n=5, *p=0.01), *domeMESO-Gal4, UAS-GFP; UAS-Ssadh^RNAi^* on RF (n=8, **p=0.001) and *domeMESO-Gal4, UAS-GFP; UAS-Ssadh^RNAi^* on SF (n=8, *p=0.02). (**b-e**) Compared to untreated **(b)** *Hml^Δ^-Gal4, UAS-GFP; UAS-Gat^RNAi^* on RF and (**c**) *Hml^Δ^-Gal4, UAS-GFP; UAS-Ssadh^RNAi^* on RF animals which show a failure of wasp-egg enwrapment, (**d, e**) 3% succinate supplementation of (**d)** *Hml^Δ^-Gal4, UAS-GFP; UAS-Gat^RNAi^* and (**e**) *Hml^Δ^-Gal4, UAS-GFP; UAS-Ssadh^RNAi^* restores it. Evident in panels **d, e** where wasp-eggs are enwrapped with lamellocytes (red). DAPI marks DNA in blue. (**f**) Progenitor (Domeless, green) maintenance or differentiation status (Peroxidasin, Pxn, red) is unaffected with succinate supplementation. Quantified in **g**, Also see Table S2. (**h**) Compared to *Hml^Δ^-Gal4, UAS-GFP; UAS-Ssadh^RNAi^* on RF (n=6), supplementing these animals with 2% fumarate (n=5) or 2% α-KG (n= 5) did not restore the lamellocyte defect.

**Supplemental Figure 4.**
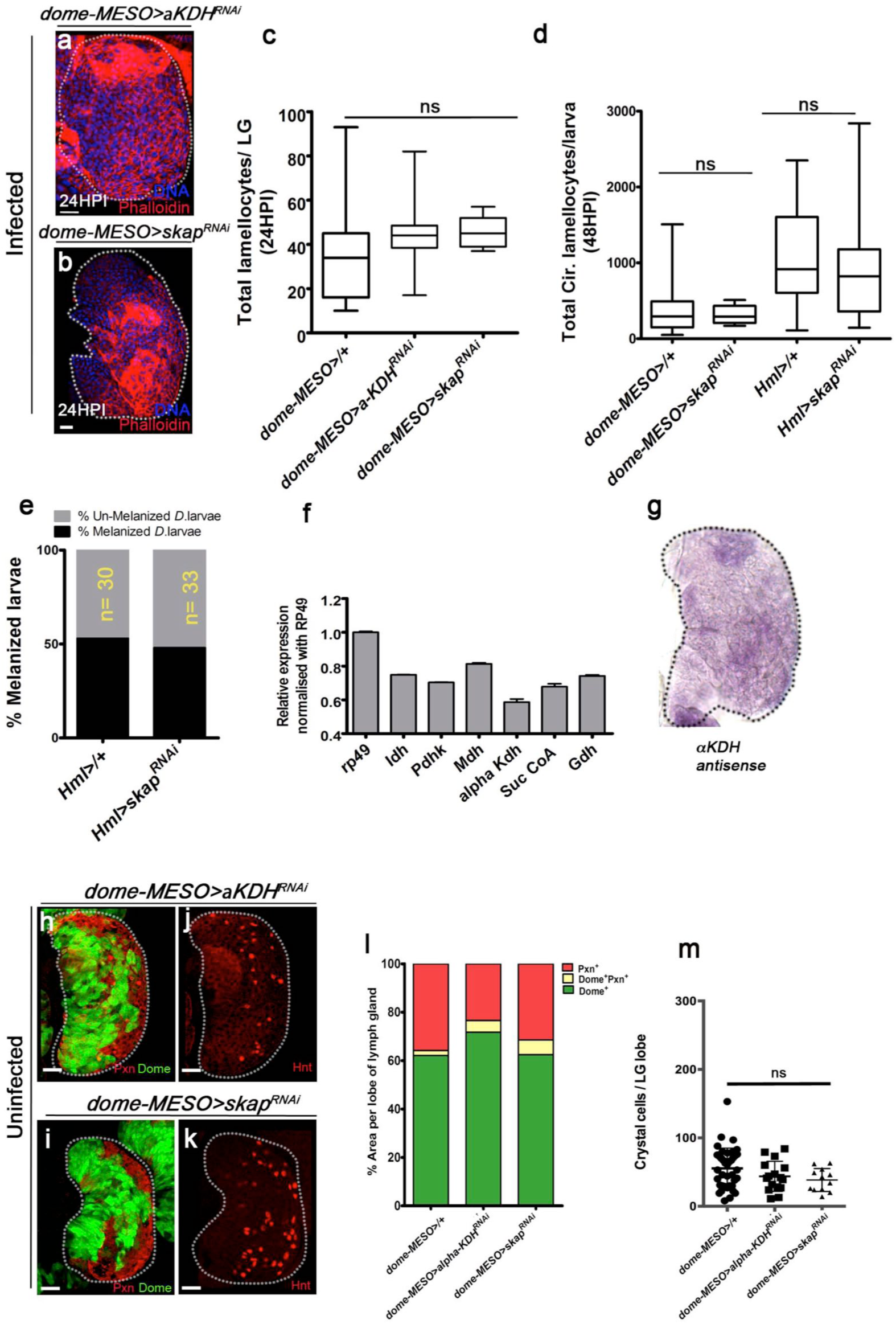
Lamellocyte induction is independent of TCA-cycle enzymes, αKDH and Skap. DNA marked with DAPI (blue), Domeless (Dome, green) positive area marks progenitor population, Peroxidasin (Pxn) marks differentiating cells and crystal cells are marked with Hindsight (Hnt). Phalloidin (red) marks blood cell outlines and lamellocytes are distinguished by their large flattened morphology. Scale bar = 20μm. HPI indicates <u>h</u>ours <u>p</u>ost wasp-infection. Bars in panel **c, d** show median in box plots and vertical bars represent upper and lowest cell counts. In **f, m** Mean ± Standard deviation (Mean ± SD) are shown. Statistical analysis in panel **c** and **d** is Mann-Whitney test, two-tailed and in **f, l, m** is un-paired *t*-test, two-tailed. “n” represents total number of larvae analyzed. (**a-c**) Progenitor-specific expression of (**a**) *α-KDH^RNAi^* (*domeMESO-Gal4, UAS-GFP; UAS-α KDH^RNAi^*) or (**b**) *skap^RNAi^* (*domeMESO-Gal4, UAS-GFP; UAS-skap^RNAi^*) does not affect lymph gland lamellocyte differentiation. (**c**) Quantifications of total lamellocytes per lymph gland lobes of *domeMESO-Gal4, UAS-GFP/+* (control, n=15), *domeMESO-Gal4, UAS-GFP; UAS-αKDH^RNAi^* (n=13) and *domeMESO-Gal4, UAS-GFP; UAS-skap^RNAi^* (n=7). (**d**) Quantifications of total circulating lamellocyte per larvae in *domeMESO-Gal4, UAS-GFP/+* (control, n*=*35), *domeMESO-Gal4, UAS-GFP; UAS-skap^RNAi^* (n=6,), *Hml^Δ^-Gal4, UAS-GFP/+* (n=15) and *Hml^Δ^-Gal4, UAS-GFP; UAS-skap^RNAi^* (n= 16) larvae. (**e**) Graphical representation of melanization response in control and *skap^RNAi^* animals (refer Table S1). (**f**) Relative quantification of *mRNA* levels of TCA-enzymes in RNA extracted from 3^rd^ instar lymph glands. The levels are normalized to *rp49* CT values. (**g**) *in situ* expression analysis of *α KDH* mRNA in 3^rd^ instar lymph glands. (**h-i**) Progenitor specific down-regulation of (**h**) *domeMESO-Gal4, UAS-GFP; UAS-α KDH^RNAi^* or (**i**) *domeMESO-Gal4, UAS-GFP; UAS-skap^RNAi^* does not affect lymph gland development, progenitor maintenance (Dome, green) and differentiation (Pxn, red). See quantifications in **l** and Table S2. (**j-k**) Down-regulation of (**j**) *domeMESO-Gal4, UAS-GFP; UAS-αKDH^RNAi^* or (**k**) *domeMESO-Gal4, UAS-GFP; UAS-skap^RNAi^* does not affect crystal cell specification (Hnt, red). See quantifications in **m** and Table S4. (**l**) Graphical representation of phenotypes described in **h** and **i**. Percentage area distribution of 3^rd^ instar larval lymph glands showing progenitor area (green, Dome^+^ only), intermediate population (IP, yellow, Dome^+^ Pxn^+^) and differentiating blood cell areas (red, Pxn^+^ only). (**m**) Crystal cell quantifications in *domeMESO-Gal4, UAS-GFP/+* (control, n=20)*, domeMESO-Gal4, UAS-GFP; UAS-αKDH^RNAi^* (n=8) and *domeMESO-Gal4, UAS-GFP; UAS-skap^RNAi^* (n=6).

**Supplemental Figure 5.**
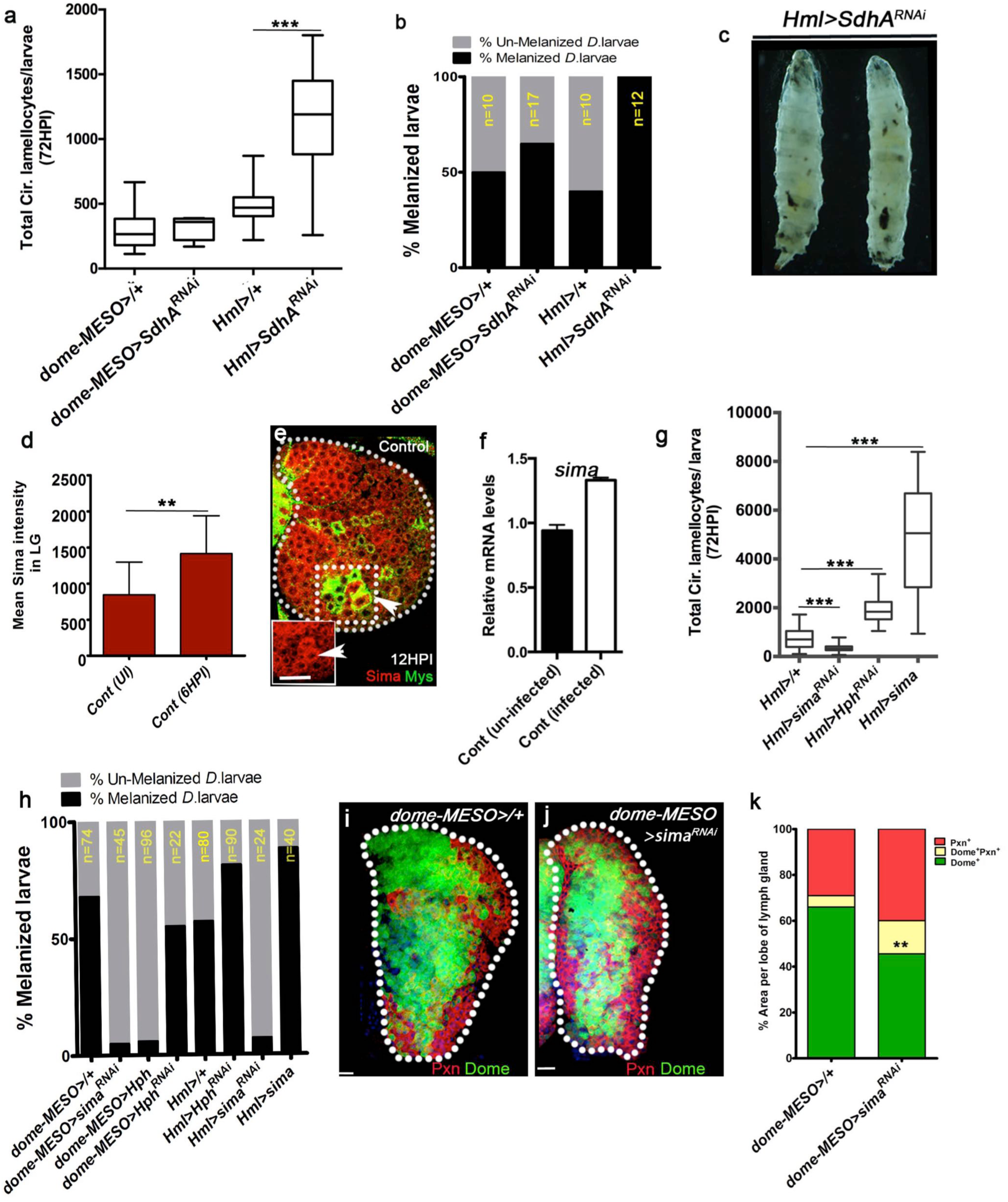
Sima function during larval hematopoiesis establishes lamellocyte potential. Dome-GFP positive area marks progenitor area. In panels **e**, **i** and **j** scale bar = 20μm. UI is un-infected, HPI indicates <u>h</u>ours <u>p</u>ost infection. In panel **a, g** median is shown in box plots and vertical bars represent upper and lowest cell counts. In **d, f, k** Mean ± Standard deviation (Mean ± SD) are shown. Statistical analysis in panel **a** and **g** is Mann-Whitney test, two-tailed and in **d, f, k** is un-paired *t*-test, two-tailed. “n” represents the total number of larvae analyzed. (**a-c**) Down-regulating *SdhA* in blood cells does not affect (**a**) lamellocyte induction or (**b, c**) melanization response. (**a**) Quantification of total circulating lamellocyte counts in *domeMESO-Gal4, UAS-GFP/+* (control, n=22), *domeMESO-Gal4, UAS-GFP; UAS-SdhA^RNAi^* (n=8), *Hml^Δ^-Gal4, UAS-GFP/+* (control, n=9), *Hml^Δ^-Gal4, UAS-GFP; UAS-SdhA^RNAi^* (n=17, ***p=0.0005). (**b**) Graphical representation of melanization response, refer Table S1 (**c**) Representative image of *Hml^Δ^-Gal4, UAS-GFP; UAS-SdhA^RNAi^ Drosophila* larvae showing enhanced melanization. (**d**) Mean intensity plot of Sima expression in lymph glands lobes from un-infected control (*domeMESO-Gal4, UAS-GFP/+*, n=18, 847 ± 451.3) and infected control at 6HPI (*domeMESO-Gal4, UAS-GFP/+*, n= 18, 1416.2 ± 523, **p=0.0013). (**e**) Control lymph glands at 12HPI showing (**e**) Sima protein expression (red) in all blood cells with comparatively higher expression detected in lamellocytes (co-stained with Myospheroid (Mys) in green and marked with white arrows in the boxed area and also shown in small inset for enhanced clarity). (**f**) Relative mRNA quantification of *sima* in control lymph glands obtained from un-infected and wasp-infected 3^rd^ instar larvae. (**g**) Quantifications of total circulating lamellocytes numbers per larvae in *Hml^Δ^-Gal4, UAS-GFP/+* (control, n=31)*, Hml^Δ^-Gal4, UAS-GFP; UAS-sima^RNAi^* (n=15, ***p=0.0003), *Hml^Δ^-Gal4, UAS-Hph^RNAi^* (n=14, ***p<0.0001) and *Hml^Δ^-Gal4, UAS-GFP/ UAS-sima* (n=30, ***p<0.0001). (**h**) Graphical representation of melanization efficiencies of the respective genotypes (Table S1). (**i, j**) Compared to (**i**) *domeMESO-Gal4, UAS-GFP/+* (control), (**j**) *domeMESO-Gal4, UAS-GFP; UAS-sima^RNAi^* expressing lymph glands show no change in progenitor (Dome, green) development, maintenance or differentiation (Pxn^+^, red). (**k**) Quantification of images shown in **i** and **j** as percentage area distribution in 3^rd^ instar larval lymph gland of progenitor (green, Dome^+^ only), intermediate population (IP, yellow, Dome^+^ Pxn^+^) and differentiating (red, Pxn^+^ only) blood cell areas. Refer Table S2.

**Supplemental Figure 6.**
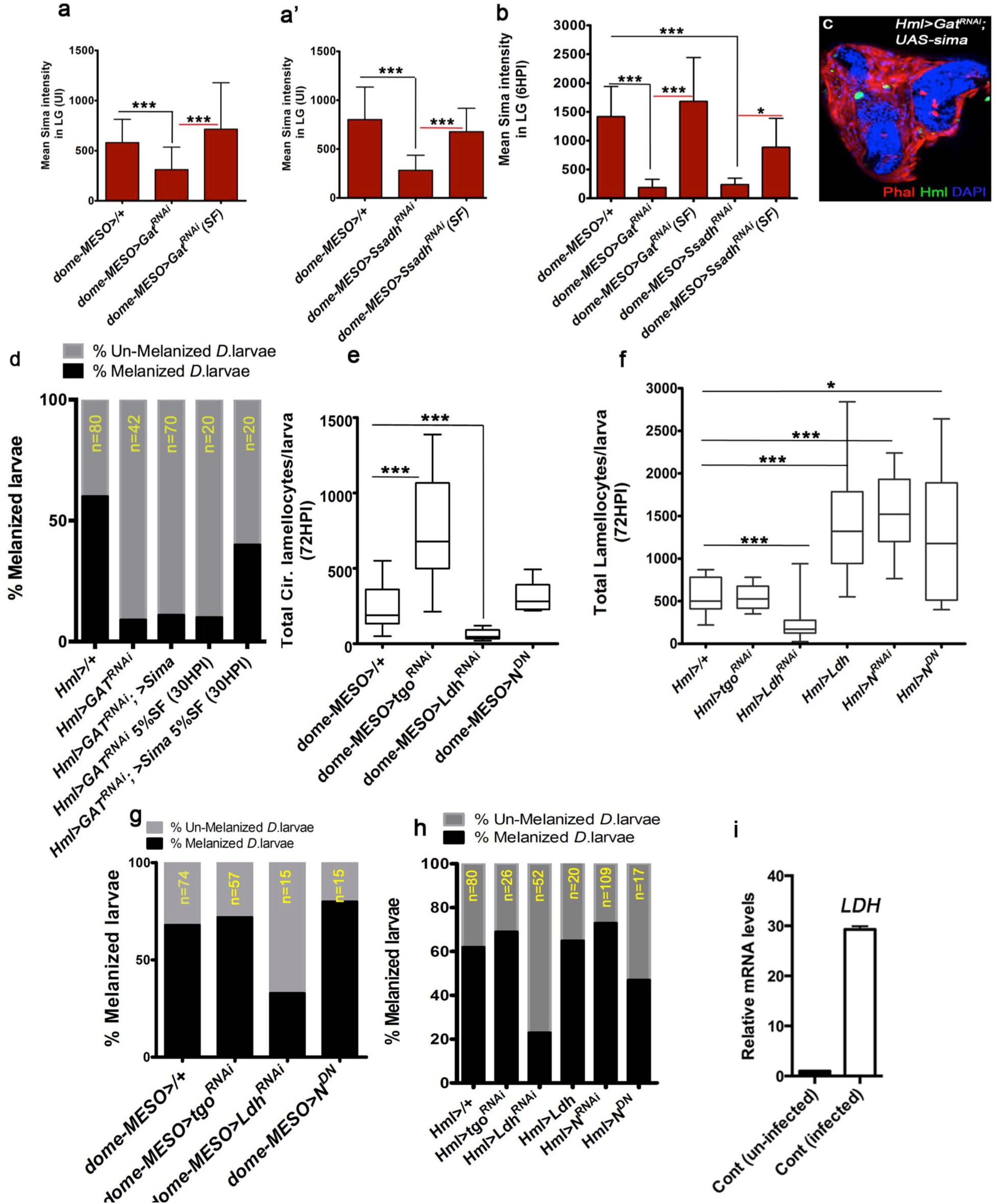
Non-canonical role for Sima in lamellocyte differentiation. UI is un-infected, HPI indicates <u>h</u>ours <u>p</u>ost infection, RF is regular food, SF is succinate food. In panels **a, b** and **i** Mean ± Standard deviation (Mean ± SD) is shown. In panels **e, f** median is shown in box plots and vertical bars represent upper and lowest cell counts. Statistical analysis in panel **a, b** and **i** is un-paired *t*-test, two-tailed and in **e, f** is Mann-Whitney test, two-tailed. “n” represents the total number of larvae analyzed. (**a, b**) Mean intensity plot of Sima expression in lymph glands lobes from (**a**) un-infected and (**b**) infected controls. (**a**) In UI, control larvae raised on RF (*domeMESO-Gal4, UAS-GFP/+*, n=25, 580.7 ± 231.6), *domeMESO-Gal4, UAS-GFP; UAS-Gat^RNAi^* on RF (n=29, 311.7 ±223.9, ***p<0.0001), *domeMESO-Gal4, UAS-GFP; UAS-Gat^RNAi^* on SF (n=16, 714.7 ± 462.5, ***p=0.0003). (**a’**) In UI, control larvae rasied on RF (*domeMESO-Gal4, UAS-GFP/+*, n=19, 801.8 ± 332.4), *domeMESO-Gal4, UAS-GFP; UAS-Ssadh^RNAi^* on RF (n=18, 282.7 ± 154.3, ***p<0.0001), *domeMESO-Gal4, UAS-GFP; UAS-Ssadh^RNAi^* on SF (n=15, 678.6 ± 238.3, ***p<0.0001). (**b**) At 6HPI, control on RF (*domeMESO-Gal4, UAS-GFP/+*, n= 18, 1416.2 ± 523, **p=0.0013 in comparison to UI control shown in **a**), *domeMESO-Gal4, UAS-GFP; UAS-Gat^RNAi^* on RF (n=18, 189 ± 141, ***p<0.0001), *domeMESO-Gal4, UAS-GFP; UAS-Gat^RNAi^* on SF (n=9, 1682 ± 760, ***p<0.0001), *domeMESO-Gal4, UAS-GFP; UAS-Ssadh^RNAi^* on RF (n=6, 238 ± 111, ***p<0.0001), *domeMESO-Gal4, UAS-GFP; UAS-Ssadh^RNAi^* on SF (n=7, 887 ± 500, *p=0.01). (**c**) Lamellocytes detected in *Hml^Δ^-Gal4, UAS-GFP/ UAS-sima; UAS-Gat^RNAi^* can encapsulate the deposited wasp-eggs (lamellocytes are marked with phalloidin in red and plasmatocytes are marked with Hml in green). (**d**) Graphical representation of melanization status, also see Table S1. (**e**, **f**) Sima function in lamellocyte specification is independent of Notch and tango function. Quantification of total circulating lamellocyte counts per larvae in (**e**) *domeMESO-Gal4, UAS-GFP/+* (control, n=27), *domeMESO-Gal4, UAS-GFP; UAS-tgo^RNAi^* (n=14, ***p<0.0001), *domeMESO-Gal4, UAS-GFP; UAS-Ldh^RNAi^* (n=14, ***p<0.0001), *domeMESO-Gal4, UAS-GFP; UAS-N^DN^* (n=6) *Hml^Δ^-Gal4, UAS-GFP/+* (control, n=29) and (**f**) *Hml^Δ^-Gal4, UAS-GFP/+* (n=11), *Hml^Δ^-Gal4, UAS-GFP; UAS-tgo^RNAi^* (n=10), *Hml^Δ^-Gal4, UAS-GFP; UAS-Ldh^RNAi^* (n=29, ***p=0.0005), *Hml^Δ^-Gal4, UAS-GFP; UAS-Ldh* (n=33, ***p<0.0001), *Hml^Δ^-Gal4, UAS-GFP; UAS-N^RNAi^* (n=10, ***p=0.0003), *Hml^Δ^-Gal4, UAS-GFP; UAS-N^DN^* (n=12, *p=0.02). (**g, h**) Graphical representation of melanization status, see Table S1. (**i**) Relative mRNA quantification of *Ldh* in control lymph glands obtained from un-infected and wasp-infected 3^rd^ instar larvae showing a 30 fold increase.

**Supplemental Figure 7.**
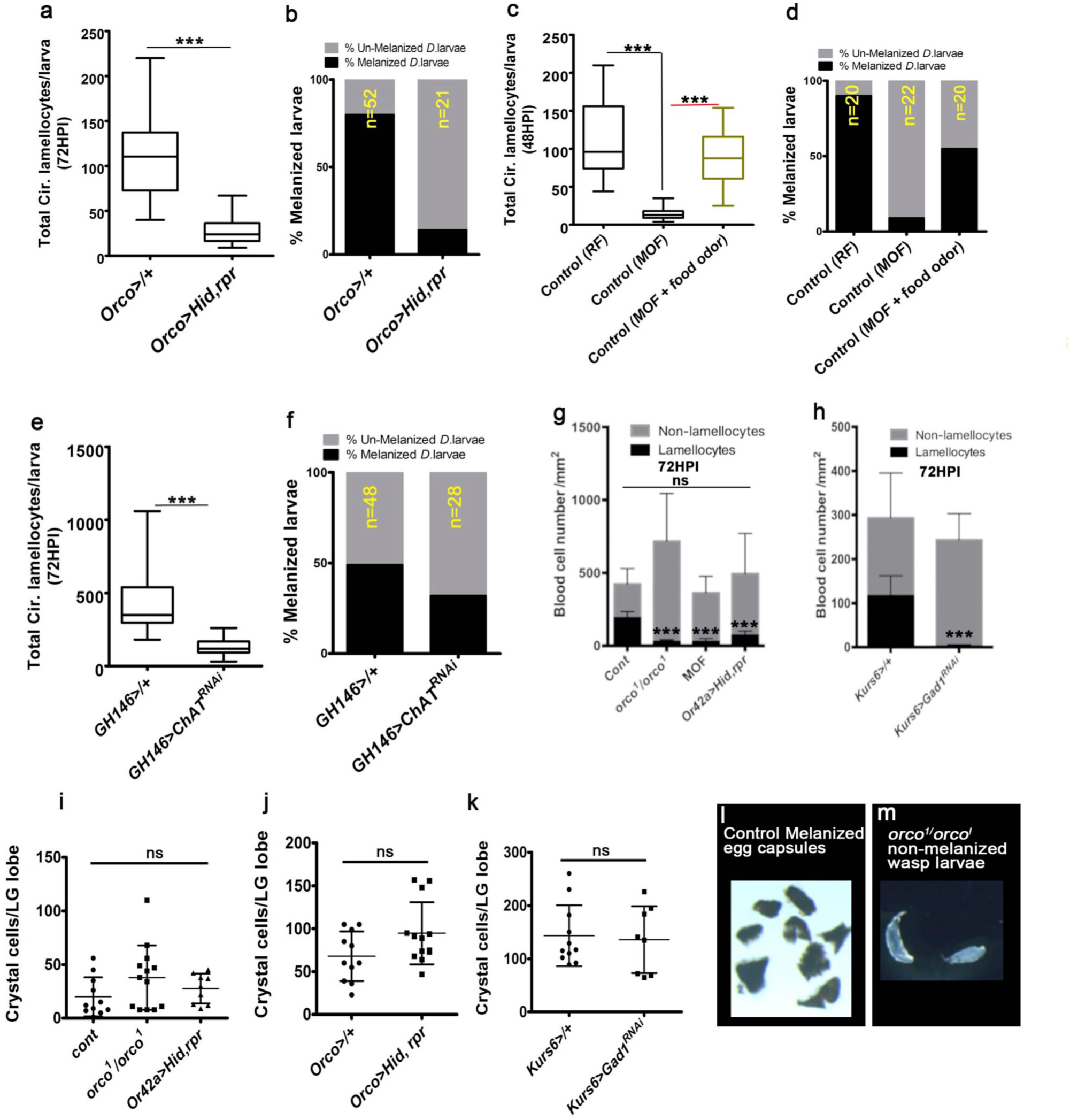
Olfaction/GABA axis controls lamellocyte induction. UI is un-infected, HPI indicates <u>h</u>ours <u>p</u>ost infection, RF is regular food, MOF is minimal odor food. In panel **a, c, e** median is shown in box plots and vertical bars represent upper and lowest cell counts. In **g-k**, Mean ± Standard deviation (Mean ± SD) are shown. Statistical analysis in panel **a, c** and **e** is Mann-Whitney test, two-tailed and in panels **g-k** is un-paired *t*-test, two-tailed. “n” represents the total number of larvae analyzed. (**a, b**) Olfactory neuron ablation inhibits lamellocyte differentiation and melanization response. Quantification of (**a**) total circulating lamellocytes in *Orco-Gal4/+* (n=34, control) and *Orco-Gal4, UAS-Hid, rpr* (n= 36, ***p<0.0001) and (**b**) corresponding melanization response (refer Table S1). (**c, d**) Larvae raised in minimal odors show lamellocyte and melanization defect which is restored with food odors. Quantification of (**c**) total circulating lamellocytes in RF (n=23), MOF (n=21, ***p<0.0001) and MOF rescue with food odors (n=18, ***p<0.0001) and (**d**) corresponding melanization in RF, MOF and MOF rescue with food odors (refer Table S1). (**e**) Blocking projection neuron signaling (*GH146>ChAT^RNAi^*) leads to reduction in lamellocyte differentiation. Quantifications of total circulating lamellocytes in *GH146>/+* (n=25, control) and *GH146>ChAT^RNAi^* (n=26, ***p<0.0001). (**d**) Quantification of melanization response seen in *GH146>/+* and *GH146>ChAT^RNAi^* animals (refer Table S1). (**g, h**) Quantification of total circulating blood cell numbers per mm^2^ in animals at 72 HPI. Represented here are the numbers of lamellocytes (black bar) and non-lamellocytes (grey bar) counted per mm^2^. No significant difference in overall cell density is detected, although a reduction in lamellocyte numbers is seen. (**g**) *orco^1^/orco^1^* (***p <0.0001), MOF (***p<0.0001), *Or42a<Hid, rpr* ((***p=0.0002) in comparison with *w^1118^*. (**h**) *Kurs6>Gad1^RNAi^* (***P=0.0001, compared to *Kurs6>/+*). Also see Table S3. (**i-k**) Quantification of total crystal cell numbers detected per lymph gland. See Table S4. (**l, m**) Melanized wasp-egg capsules detected in (**l**) control animals is (**m**) not seen in *orco^1^/orco^1^* mutant larvae.

**Supplemental Figure 8.**
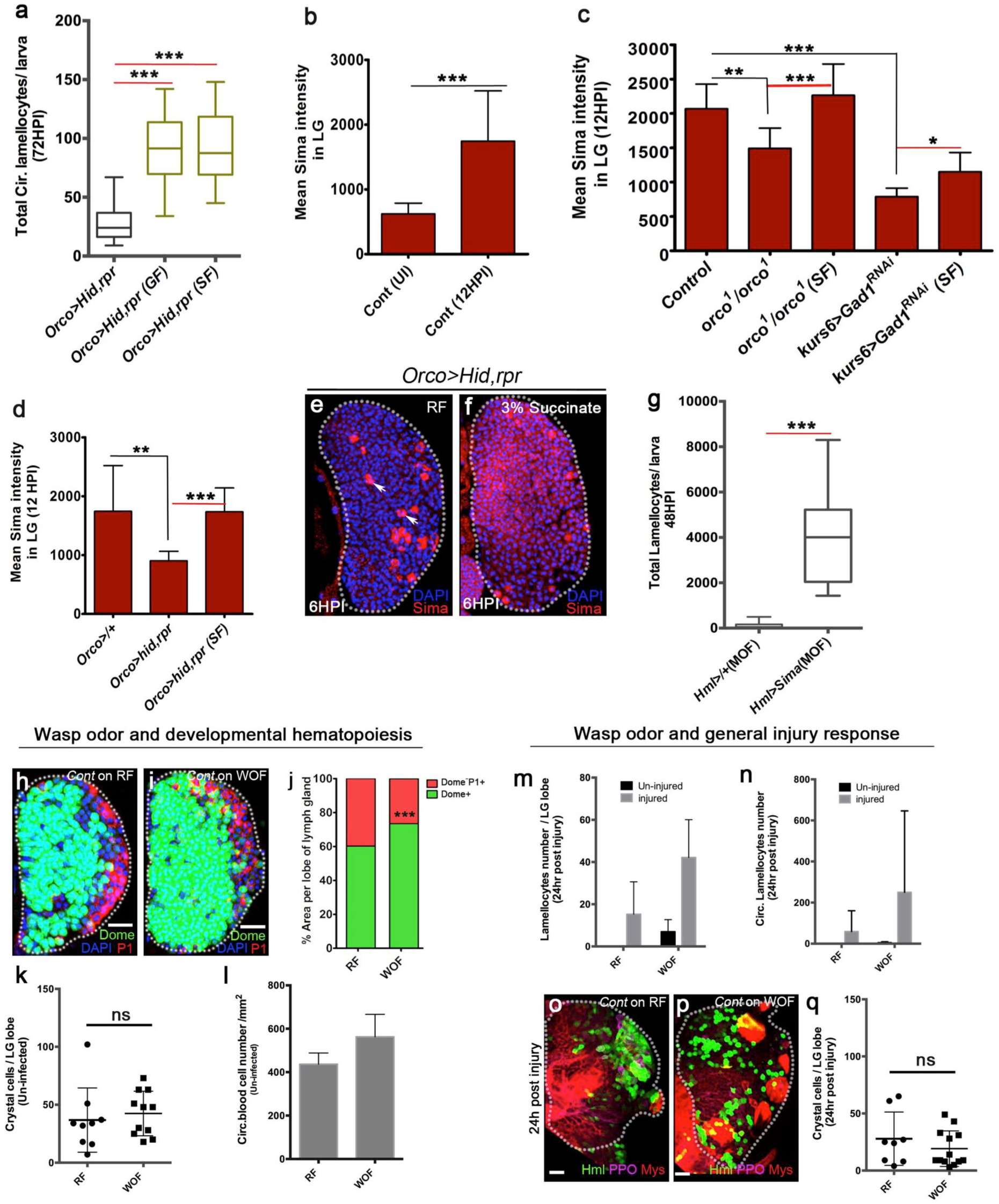
Olfaction/GABA axis controls lamellocyte induction via modulating blood cell succinate and Sima levels. Nuclei are marked with DAPI (blue), Domeless (Dome, green) marks progenitors, P1 (red) marks plasmatocytes, Hemolectin (Hml, green) marks differentiating blood cells, prophenol oxidase (PPO, magenta) marks crystal cells; Myospheroid (Mys, red) marks lamellocytes. RF is regular food, GF is GABA food, SF is succinate food, MOF is minimal odor food, WOF is wasp odor food, ns is non-significant. In **a, g** median is shown in box plots and vertical bars represent upper and lowest cell counts. In **b-d, j-n** and **q** Mean ± Standard deviation (Mean ± SD) are shown. Statistical analysis in panel **a, g** is Mann-Whitney test, two-tailed and in panels **b-d, j-n** and **q** is un-paired *t*-test, two-tailed. “n” represents the total number of larvae analyzed. Scale bars = 20μm. (**a**) Quantifications showing total circulating lamellocyte counts per larvae in *Orco-Gal4, UAS-Hid, rpr* on RF (n= 36) and its rescue on GF (n=16, ***p<0.0001) and succinate food (n=20, ***p<0.0001). (**b**) Mean intensity plot of Sima expression in control (*Orco>/+*) lymph glands lobes in un-infected (n= 20, 622 ± 166) and infected (n=12, 1745 ± 775.2, ***p<0.0001) states. (**c**) Mean intensity plot of Sima expression in lymph glands lobes at 12HPI in (**c**) control on RF (*w^1118^*, n=9, 2069 ± 359), *orco^1^/orco^1^* on RF (n=9, 1493 ± 293, **p= 0.0018), *orco^1^/orco^1^* on SF (n=10, 2266 ± 455,, ***p=0.0004), *Kurs6>Gad1^RNAi^* on RF (n=6, 789.4 ± 124, ***p<0.0001), *Kurs6>Gad1^RNAi^* on SF (n=8, 1151 ± 281, *p=0.0126). Representative images shown in **Fig. 6a-e**. (**d-f**) Lymph glands of infected *Orco-Gal4, UAS-Hid, rpr* animals show reduced Sima protein expression. (**d**) Mean intensity plot of Sima expression in lymph glands lobes at 12HPI of *Orco>/+* on RF (n=12, 1745 ± 775), *Orco>UAS-hid, rpr* on RF (n=11, 903.7 ± 161, **p=0.002), *Orco>UAS-hid, rpr* on SF (n=12, 1736 ± 405, ***p<0.0001). (**e, f**) Representative lymph gland images of infected *Orco-Gal4, UAS-Hid, rpr* animals showing (**e**) reduced Sima protein expression (red) in blood cells except in crystal cells (white arrows) (**f**) which is restored with succinate. (**g**) Quantifications showing total circulating lamellocyte counts in animals raised on MOF (n= 12) restored by blood-cell specific Sima expression (*Hml^Δ^-Gal4, UAS-GFP; UAS-sima*, n=13, ***p<0.0001). (**h-j**) Representative 3^rd^ instar lymph gland images from *domeMESO>GFP/+* showing progenitors (Domeless, green) and plasmatocytes (red) in (**h**) RF and (**i**) WOF and quantifications of their respective areas in **j (**progenitor area in green and differentiated blood cell area in red). Also see Table S2. (**k**) Quantification of crystal cell numbers per lymph gland lobe in uninfected animals from RF (n=5) and WOF (n=6). Also see Table S4. (**l**) Quantification of circulating blood cell numbers/mm^2^ in uninfected animals (*domeMESO>GFP/+*) from RF and WOF (for each condition n=5). (**m, n**) Quantification of lamellocyte numbers in *Hml^Δ^>GFP/+* (**m**) lymph glands and (**n**) circulation in un-injured (black bar, control) and 24 hours post-injury animals (grey bar). (**o, p**) Differentiating blood cells (green), lamellocytes (red) and crystal cells (magenta) marked in lymph glands of (**o**) controls (RF) and (**p**) 24 hours post-injury animals (*Hml^Δ^>GFP/+*) raised in WOF. (**q**) Quantification of crystal cell numbers in lymph glands (*Hml^Δ^>GFP/+*) at 24 hours post injury of RF (n=4) and WOF (n=7) larvae.

**Supplemental Figure 9.**
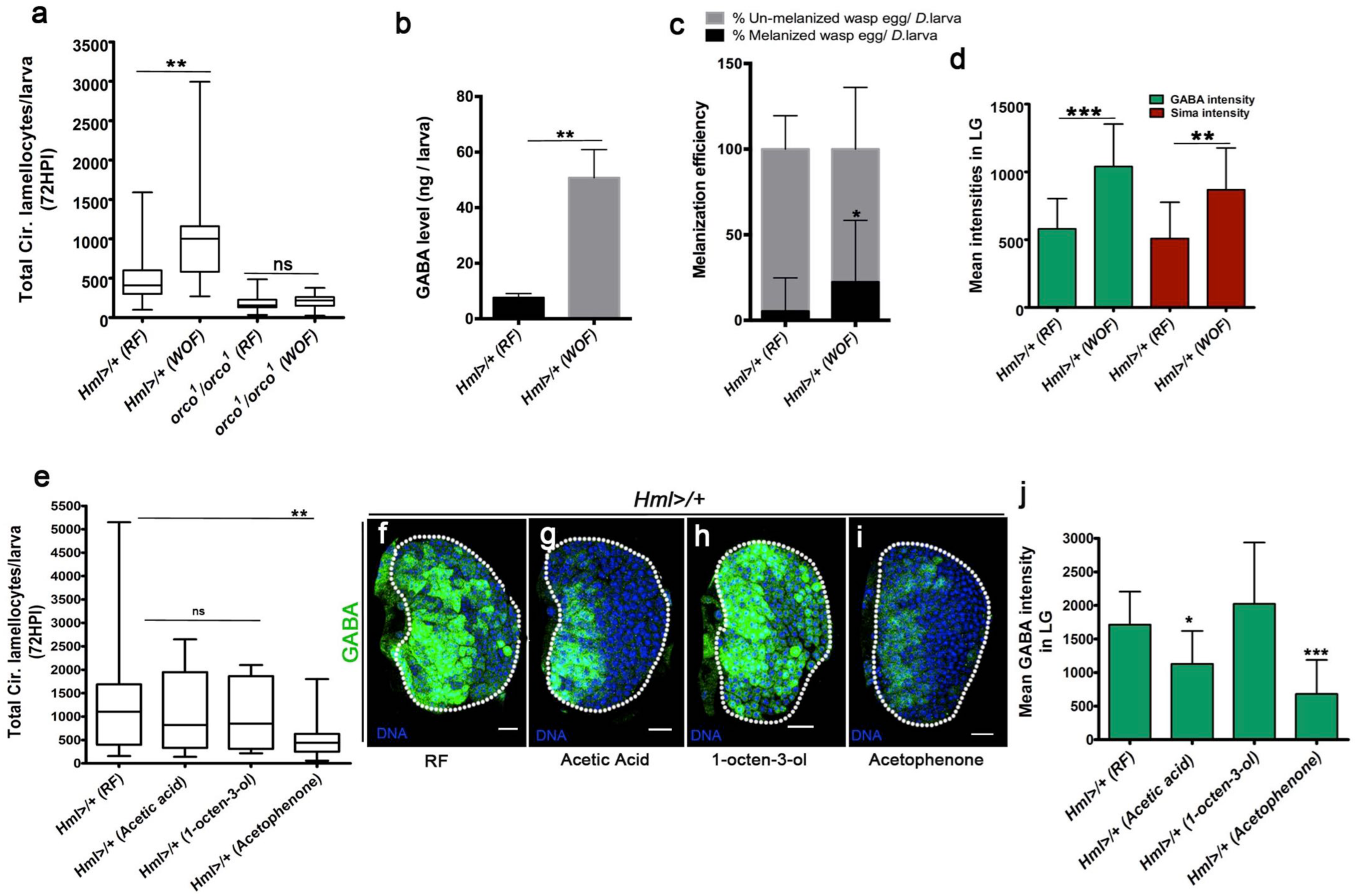
Physiological control of cellular immunity by pathogenic wasp-odors. Representative 3rd instar lymph gland images (**f-i**) showing DNA stained with DAPI (blue) and intra-cellular GABA (iGABA, green). Scale bar = 20mm. HPI indicates <u>h</u>ours <u>p</u>ost wasp-infection, RF is regular food, WOF is wasp odor food and ns is non-significant. In **b, c, d, j** Mean ± Standard deviation (Mean ± SD) is shown. In **a** and **e** median is shown in box plots and vertical bars represent upper and lowest cell counts. Statistical analysis in panels **b, c, d, j** is un-paired *t*-test, two-tailed and in **a** and **e** is Mann-Whitney, two-tailed. “n” represents the total number of larvae analyzed. (**a**) Total circulating lamellocyte quantification in RF (*Hml>/+*, n=19) and WOF (*Hml>/+*, n=19, **p=0.009), *orco^1^/orco^1^* on RF (n=27) and *orco^1^/orco^1^* on WOF (n=25). (**b**) Compared to hemolymph GABA levels in larvae raised on RF, WOF larvae have elevated systemic GABA. Quantifications of hemolymph GABA from 3^rd^ instar larvae on RF (*Hml>/+*), WOF (*Hml>/+*, **p=0.002). Refer Table S5 for absolute amounts. (**c**) Quantification of increased melanization efficiencies in WOF animals (*Hml>/+*, n=32, *p=0.02) in comparison to controls on RF (*Hml>/+*, n=33). (**d**) Mean intensity plot of lymph gland lobes iGABA (green bars) and Sima protein (red bars) expression. *Hml^Δ^>GFP/+* on RF (iGABA, n=18, 579 ± 224; Sima, n=12, 509 ± 267), and *Hml^Δ^>GFP/+* on WOF (iGABA, n=9, 1040 ± 313, ***p=0.0002; Sima, n=12, 869 ± 309, **p=0.006). (**e**) Quantification of total circulating lamellocyte numbers per larvae in controls (*Hml>UAS-GFP/+*) when raised on RF (n=49), acetic acid (n=11), 1-octen-3-ol (n=10) and acetophenone (n=23, **p=0.0013). (**f-j**) iGABA levels in *Hml>UAS-GFP/+ 3^rd^* instar lymph glands obtained from animals reared in different odors conditions. Compared to iGABA detected in (**f**) RF (*Hml>UAS-GFP/+*) **(g)** acetic acid shows some reduction, (**h**) 1-octen-3-ol shows no change, and **(i)** acetophenone leads to significant reduction. **(j)** Mean intensity plots of lymph glands lobes iGABA (green bars). *Hml^Δ^>GFP/+* on RF (iGABA, n=17, 1713.6 ± 492.6), *Hml^Δ^>GFP/+* on acetic acid (iGABA, n=8, 1128.8 ± 492.1, *p=0.0109), 1-octen-3-ol (iGABA, n=11, 2023.6 ± 914.3) and acetophenone (iGABA, n=10, 680.8 ± 508.0, ***p<0.0001).

**Supplemental Figure 10.**
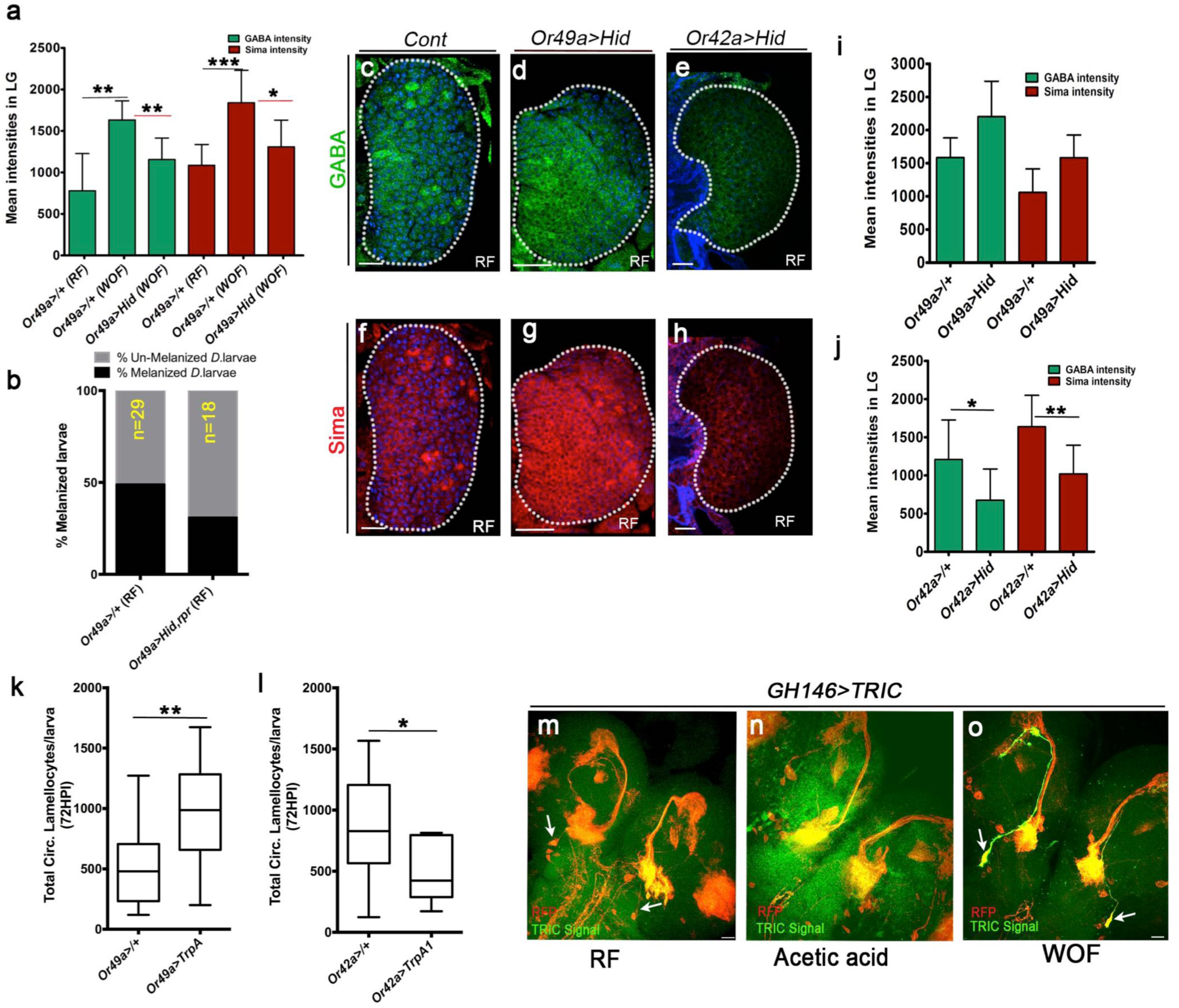
Specific wasp ORN activation drives lamellocyte expansion. DNA marked with DAPI (blue), iGABA (green), Sima (red). Scale bars = 20μm. HPI indicates <u>h</u>ours <u>p</u>ost wasp-infection, RF is regular food, WOF is wasp odor food and ns is non-significant. In **a, i** and **j** Mean ± Standard deviation (Mean ± SD) is shown. In **k** and **l** median is shown in box plots and vertical bars represent upper and lowest cell counts. Statistical analysis in panels **a, i** and **j** is un-paired *t*-test, two-tailed and in **k** and **l** is Mann-Whitney, two-tailed. “n” represents the total number of larvae analyzed. (**a**) *Or49a>/+* on RF (iGABA, n=8, 779 ± 448; Sima, n=8, 1085 ± 251.4), *Or49a>/+* on WOF (iGABA, n=6, 1632 ± 230, **p=0.0012; Sima n=8, 1840 ± 390, ***p=0.0004), *Or49a-Gal4, UAS-Hid* on WOF (iGABA, n=8, 1156 ± 256.4, **p=0.004; Sima, n=8, 1308 ± 323, *p=0.01). Representative images shown in main Figure 7f, h and **j,k**. (**b**) Quantification of melanization response in *Or49a-Gal4; UAS-Hid* animals, see Table S1. (**c-h**) Unlike Or42a, Or49a function is dispensable for lymph gland iGABA and Sima protein expression. Shown in (**c-e**) iGABA (green) and (f-h) Sima (red) expression. Compared to (**c, f**) control (*Or49a>/+*), (**d, g**) *Or49a-Gal4, UAS-Hid* lymph glands show no difference in expression. (**e, h**) iGABA and Sima levels are reduced in *Or42a-gal4, UAS-Hid* animals. Quantified in **i, j**. (**i, j**) Mean intensity plots of lymph glands lobes iGABA (green bars) and Sima protein (red bars) expression upon blocking (**i**) Or49a or (**j**) Or42a function. (**i**) GABA, *Or49a>/+* (control, n= 10, 1585± 295), *Or49a>UAS-Hid* (n=5, 2201 ± 535, p=0.01) and Sima, *Or49a>/+* (control, n= 10, 1059 ± 356), *Or49a>UAS-Hid* (n=5, 1583 ± 340, p=0.02). (**j**) GABA, *Or42a>/+* (control, n= 10, 1209 ± 517), *Or42a>UAS-Hid* (n=10, 676 ± 408, *p=0.02) and Sima in *Or42a>/+* (control, n= 10, 1638 ± 411), *Or42a>UAS-Hid* (n=10, 1021 ± 376, **p=0.002). (**k, l**) Quantifications of total circulating lamellocytes seen upon forced activation of (**k**) *Or49a>/+* (control, n=19) and *Or49a-Gal4, UAS-TrpA1* (n=17, **p=0.001) and (**l**) *Or42a>/+* (control, n=17), *Or42a-Gal4, UAS-TrpA1* (n= 10, *p=0.03). (**m-o**) Differential control of odor exposure on PN activity. PN activity was assessed by monitoring intracellular calcium signaling using the transcriptional reporter (TRIC). TRIC is designed to detect changes in neuronal activity where green shown Ca^2+^ activity and red (*UAS-RFP*) marks the neurons expressing *GH146>TRIC*. Compared to TRIC activity (GFP) detected in 3^rd^ instar larval brain tissue (**m**) from RF and (**n**) acetic acid, (**o**) WOF condition shows elevated Ca^2+^ reporter activity in a specific subset of PN-neurons (marked by white arrows in **o**). These same neurons in RF and acetic acid (red, and marked by white arrows) do not show any TRIC activity. Implying long-term effect of wasp odors on PN-firing.

**Supplemental Figure 11.**
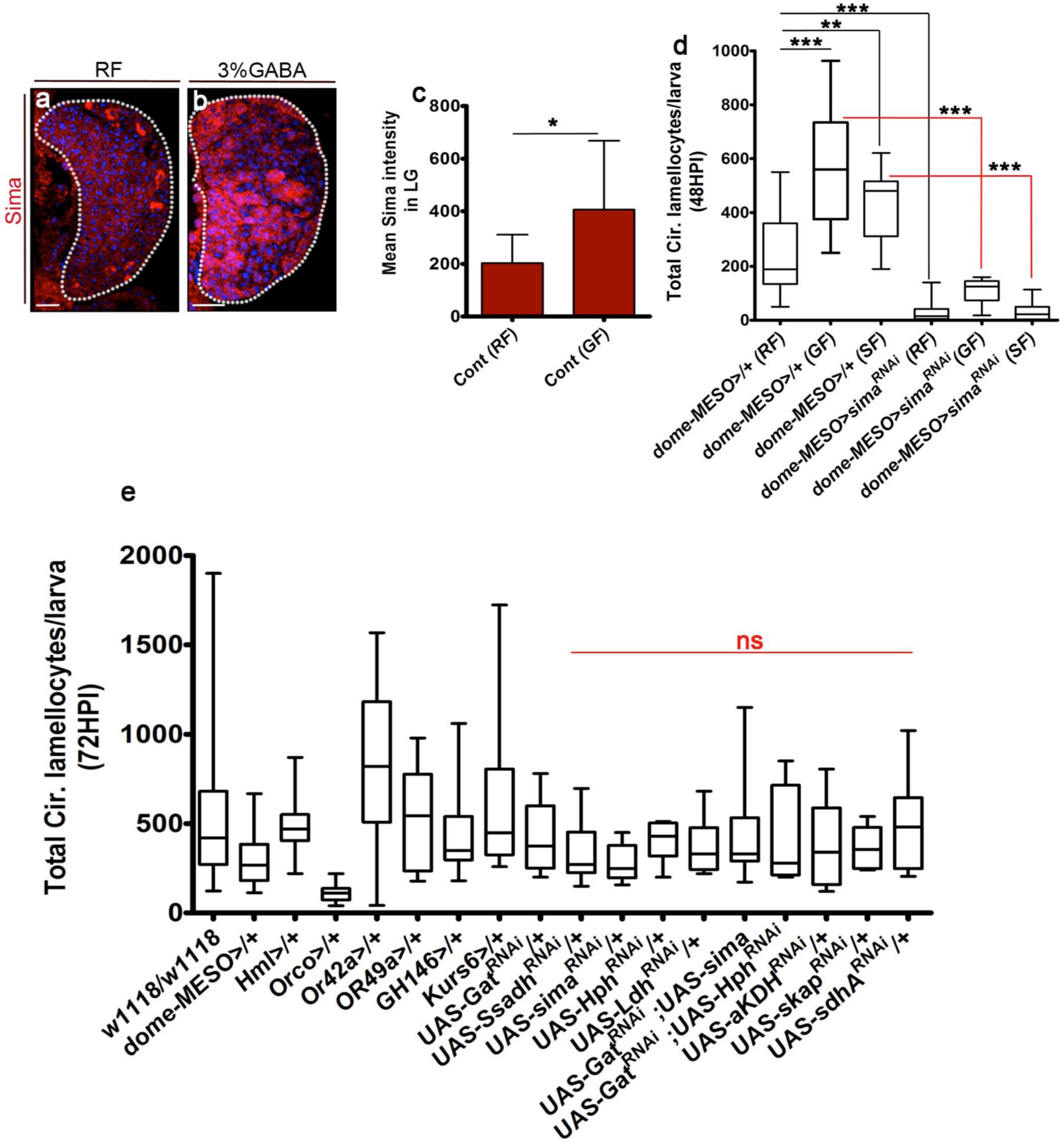
GABA availability regulates lamellocyte potential. DNA marked with DAPI (blue). Scale bar = 20μm. RF indicates regular food, GF is GABA food, HPI indicates <u>h</u>ours <u>p</u>ost wasp-infection. In **c** Mean ± Standard deviation (Mean ± SD) are shown. In **d, e** median is shown in box plots and vertical bars represent upper and lowest cell counts. Statistical analysis in panels **c** is un-paired *t*-test, two-tailed and **d, e** is Mann-Whitney test, two-tailed. “n” represents the total number of larvae analyzed. Scale bars = 20μm. (**a-c**) Comparative expression analysis of Sima protein (red) in lymph gland in animals raised on (**a**) RF and (**b**) GF. Elevated Sima is seen in GABA fed animals. (**c**) Corresponding quantifications of mean Sima intensities in the lymph glands lobes on RF (control, n=11, 203 ± 108) and GF (control, n= 8, 406 ± 262 *p=0.03). (**d**) Quantification of total circulating lamellocyte numbers per larvae in *domeMESO-Gal4, UAS-GFP/+* on RF (control, n=27), 3% GABA (n=9, ***p=0.0003) and 3% succinate (n=7, **p=0.008). *domeMESO-Gal4, UAS-GFP; UAS-sima^RNAi^* (n=29, ***p <0.0001) abrogates lamellocyte expansion seen in 3% GABA (n=9, ***p<0.0001) and 3% succinate (n=24, ***p<0.0001). (**e**) Quantification of total circulating lamellocytes numbers in all the *Gal4* lines used in this study, *RNAi* constructs without the *Gal4* and the genetic rescue constructs. All the lines have been crossed-out to *w^1118^* (not significant in comparison to *w^1118^*).

**Table S1.**
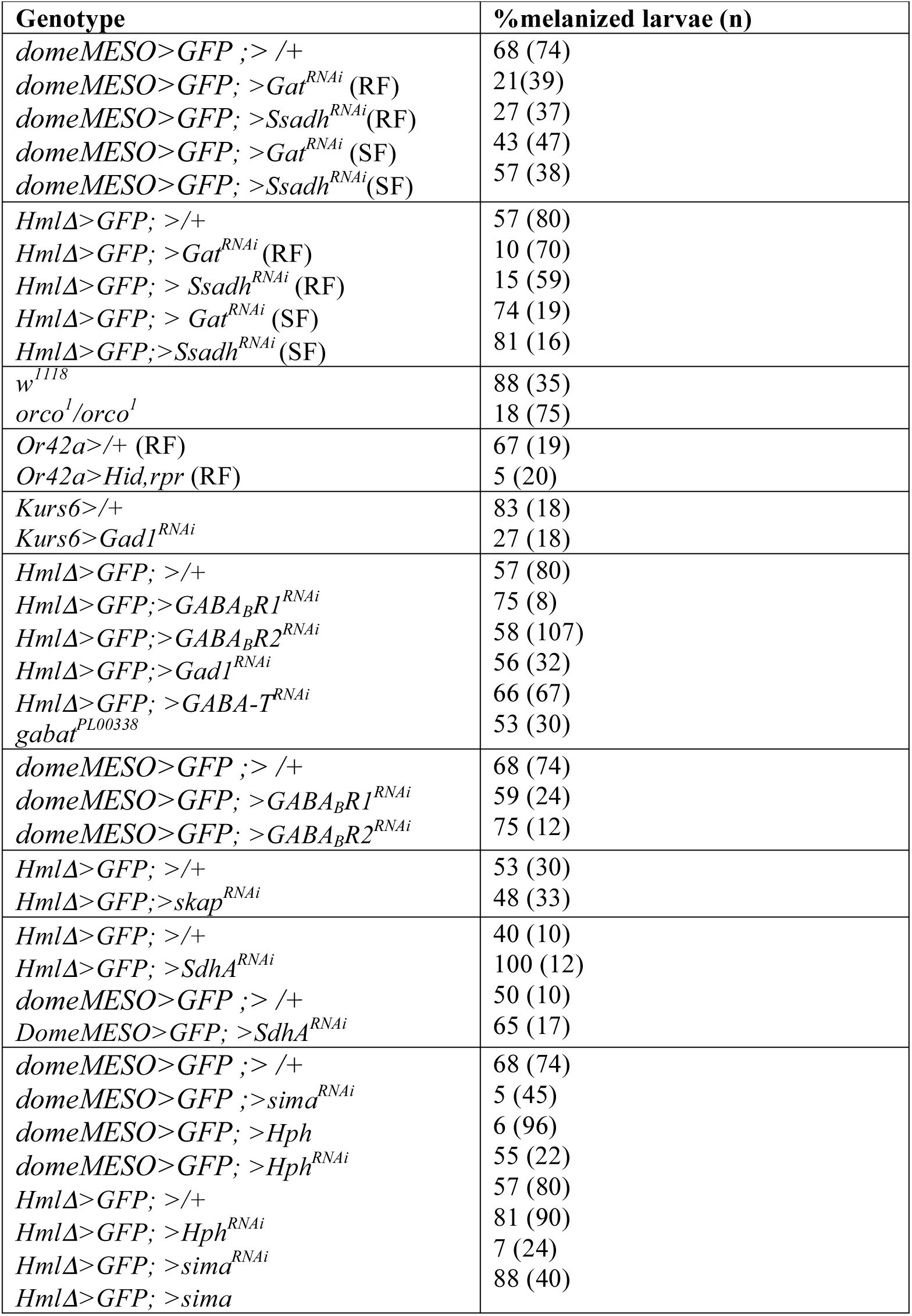
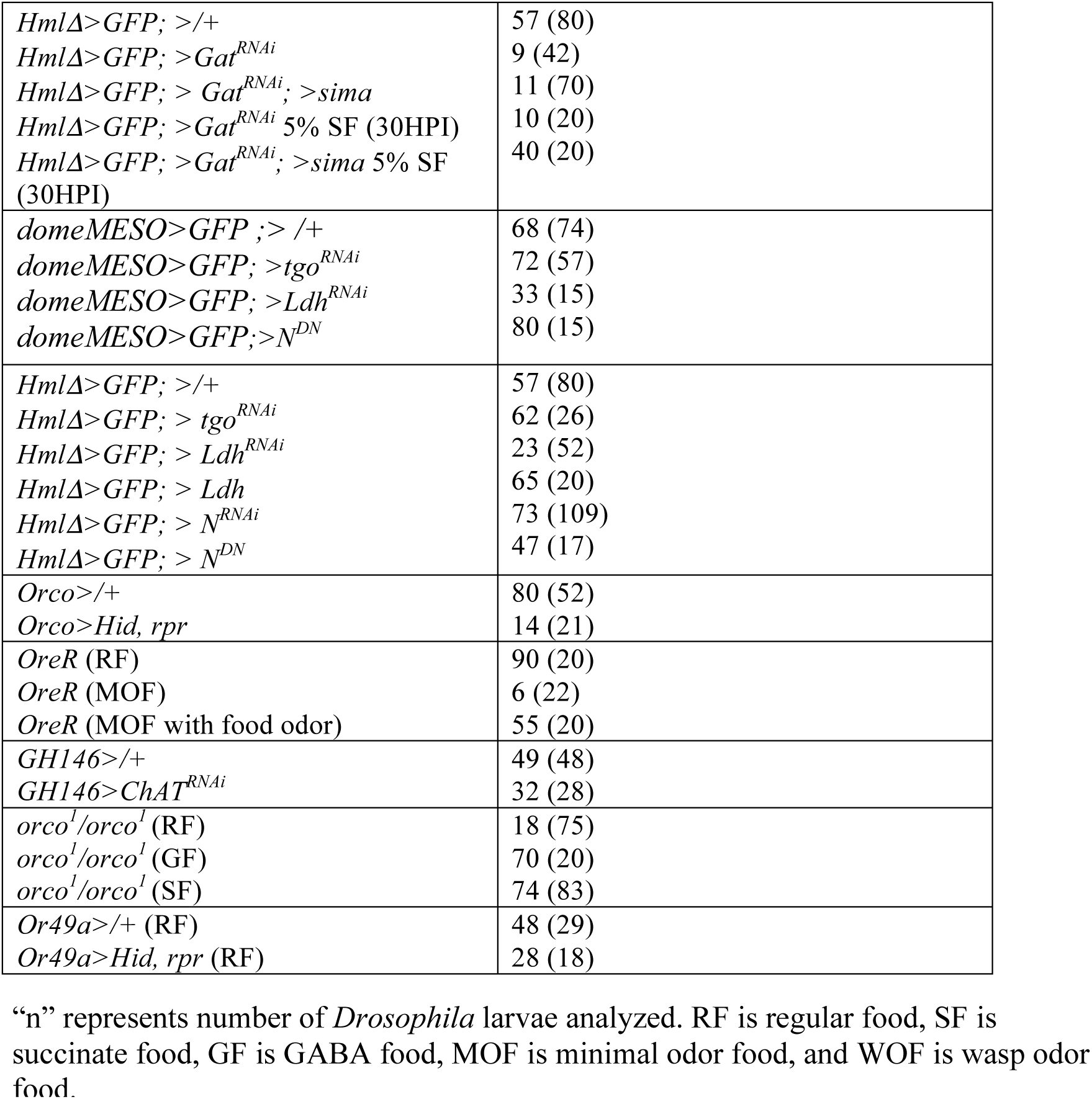
Quantification of melanization status.

**Table S2.**
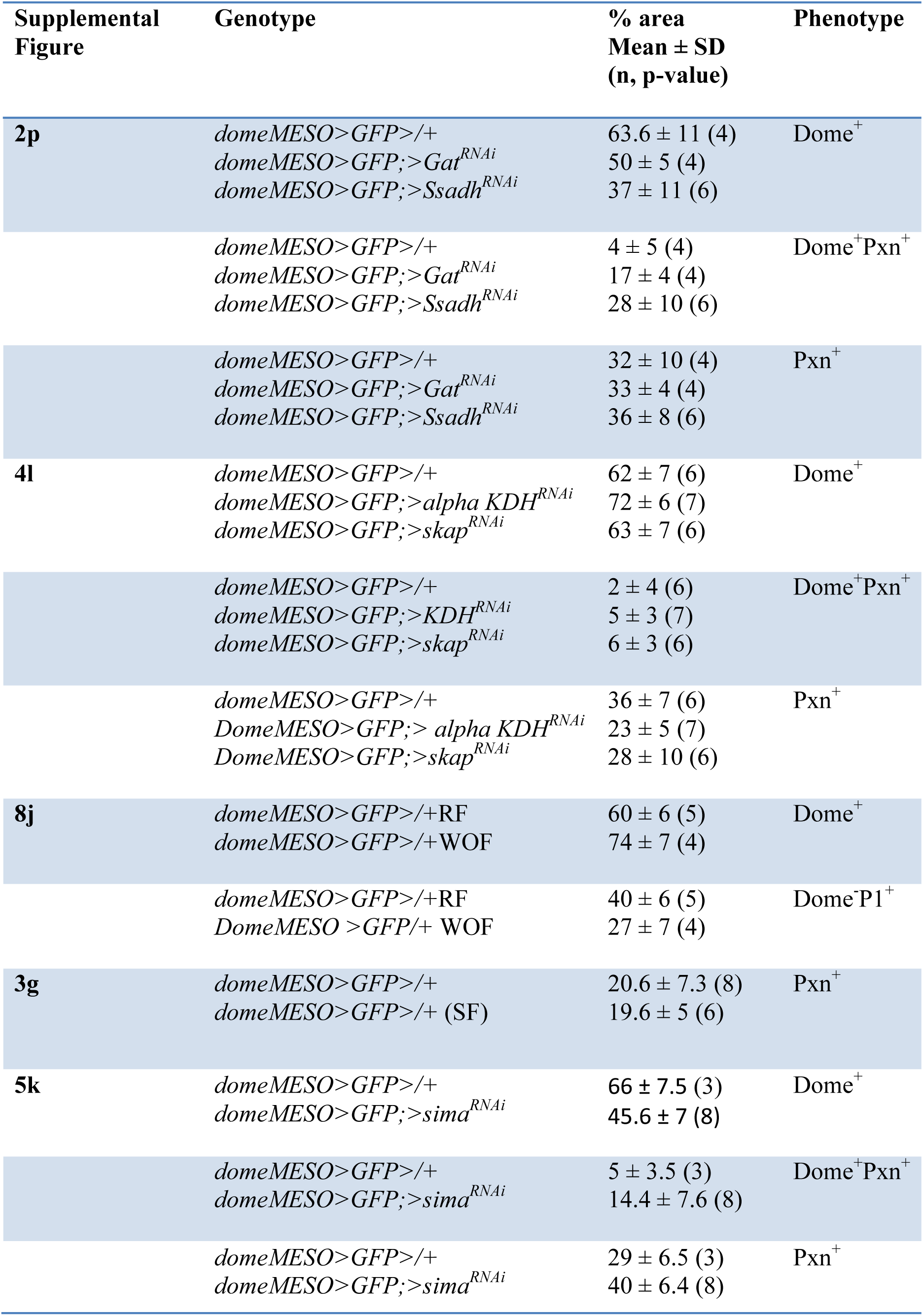
Lymph gland area quantifications.

**Table S3.**
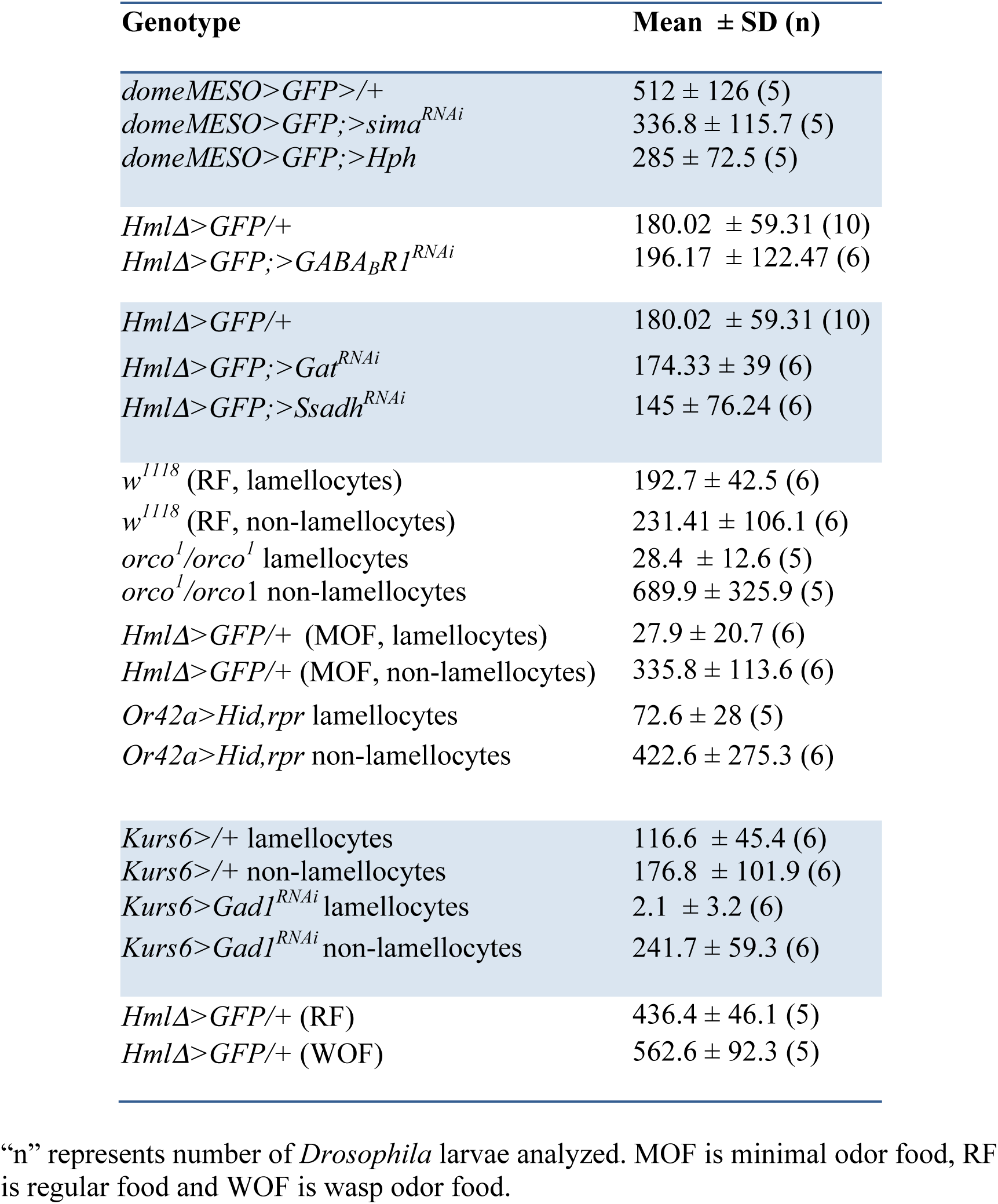
Circulating cell counts represented as cells/mm^2^.

**Table S4.**
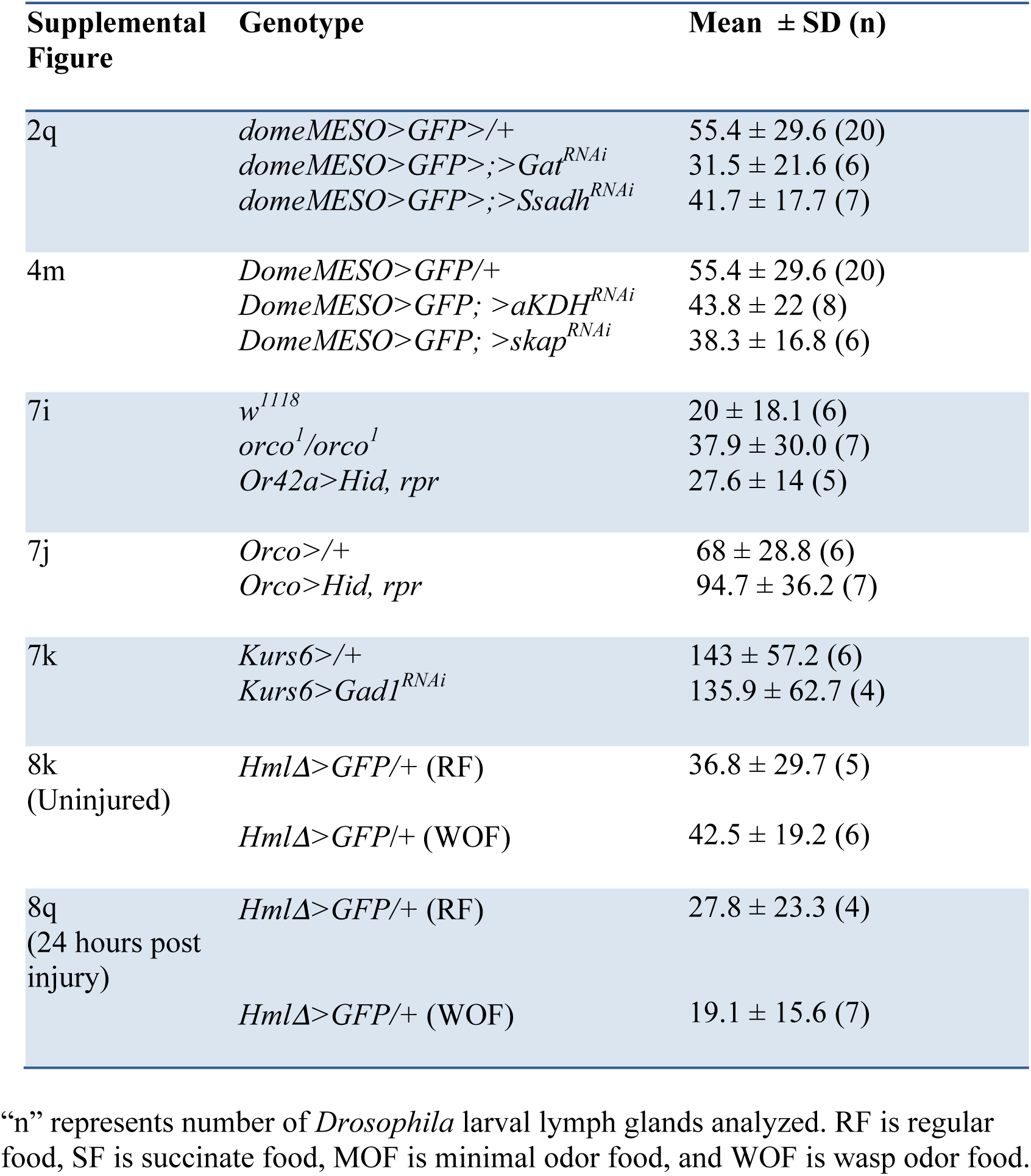
Crystal cell counts per lymph gland lobe.

**Table S5.** Hemolymph GABA measurement.

| **Figure** | **Genotype** | **GABA (ng/larvae)**<br>**Mean ± SD (n, N)** | **p-value** |
| --- | --- | --- | --- |
| Fig 7b | *Or49a*>/+ (RF) | 46 ± 25.8 (3, 9) | |
| | *Or49a*>/+ (WOF) | 100.5 ± 20.3 (3, 9) | 0.0008 (WOF vs RF) |
| | *Or49a*>*Hid, rpr* (WOF) | 63.7 ± 20.3 (5, 6) | 0.0105 (vs WOF) |
| | *Or49a*>*Hid, rpr* (RF) | 60.7 ± 27 (5, 6) | 0.25 (vs RF) |
| Supplemental Fig 9b | *Hml*Δ>/+ (RF) | 7.6 ± 1.5 (5, 3) | |
| | *Hml*Δ>/+ (WOF) | 50.8 ± 1.5 (5, 3) | 0.002 (WOF vs RF) |

“n” represents total number of *Drosophila* larvae bled to obtain hemolymph for GABA measurement and “N” represents the number of biological repeats. RF is regular food, SF is succinate food, MOF is minimal odor food and WOF is wasp odor food.

